# ERK-dependent DICER1 phosphorylation promotes open chromatin state and lineage plasticity to mediate tumor progression

**DOI:** 10.1101/2022.11.01.514714

**Authors:** Raisa A. Reyes-Castro, Shin-Yu Chen, Jacob Seemann, Swathi Arur

**Affiliations:** The University of Texas MD Anderson Cancer Center. Department of Genetics. Houston Texas; University of Puerto Rico, School of Medicine, San Juan, Puerto Rico; The University of Texas MD Anderson Cancer Center UTHealth Houston Graduate School of Biomedical Sciences., Genetics and Epigenetic program, Houston, Texas

## Abstract

DICER1 controls micro(mi)RNA-mediated epithelial-to-mesenchymal transition (EMT) to regulate tumorigenesis of lung adenocarcinomas (LUADs). We discovered that DICER1 is phosphorylated by ERK and nuclear translocated and phospho-DICER1 contributes to tumorigenesis. Mechanisms through which phospho-DICER1 regulates tumor progression remain undefined. We show that phospho-nuclear DICER1 associates with invasive human LUADs with oncogenic *KRAS* mutations and promotes late-stage tumor progression in mice with oncogenic *Kras* mutations. Surprisingly, phosphomimetic DICER1 regulates LUAD progression independent of miRNAs and EMT. Integrating single-cell RNA sequencing, fluorescent *in situ* RNA hybridization, immunofluorescence, and ATAC-sequencing, in mice, we discovered that phosphomimetic DICER1 generates an open chromatin state in the lung tumor alveolar cells leading to expression of gastrointestinal genes and altered AT2 cell identity. Strikingly, we also observe the gastric gene signature in human LUADs with phospho-DICER1 and *KRAS* mutations. We propose that phosphorylated nuclear DICER1 regulates chromatin remodeling leading to tumor cell reprogramming which drives lung cancer progression.

## Introduction

Lung cancer is the leading cause of cancer-related mortality worldwide primarily due to the formation of metastatic lesions, which induce multi-organ failure^1^. Oncogenic gain-of-function mutations in the small GTPase *KRAS* account for 20-30% of human lung adenocarcinomas (LUADs)^2-5^. The oncogenic mutations in the GTPase domain render KRAS locked in the GTP-loaded and thus ‘active’ state^6, 7^ Notably, expression of oncogenic mutant *Kras* is sufficient to cause cell transformation and lung tumor formation in murine models^8^. Molecularly, KRAS functions through the activation of the core kinase cascade RAF-MEK-ERK (Extracellular signal Regulated Kinase) to control a multitude of cellular behaviors^9^. More specifically, active di-phosphorylated ERK which is the final effector of the cascade phosphorylates diverse substrates that in turn control cellular processes such as proliferation, migration, apoptosis, and differentiation, among others^10, 11^. Thus, to understand the pathobiology of lung adenocarcinomas driven by the KRAS-ERK signaling pathway we need to identify and define the substrates through which *KRAS* oncogenic mutations specifically drive tumor onset, progression, and invasion. However, unique targets of ERK that mediate only late stages of cancer progression and invasion remain undefined.

We discovered DICER1 as a unique substrate of active ERK in worms, mice and humans^12-14^. ERK phosphorylates DICER1 at two serine residues in the RNaseIIIb domain and the dsRNA Binding Domain (RBD)^12-15^. In cancers, *Dicer1* functions as a haploinsufficient tumor suppressor^16, 17^. In this context, loss of one copy of *Dicer1* in a *Kras* oncogenic murine background leads to an increase in tumor number and size. Yet, somatic mutations that result in deletion or loss of *Dicer1* are rare in cancers^18-20^. Thus, the role of *Dicer1* in cancer onset and progression is nuanced and indicates that *Dicer1* function is context dependent. We hypothesized that phosphorylation of DICER1 likely provides one context for regulation of DICER1 activity and tested this function in cancer development. Genetically modified mouse models that carry the constitutive homozygous form of phosphorylated DICER1 (phosphomimetic *Dicer1; Dicer1*^S2D^) wherein the phosphorylated serines at positions 1712 and 1836 are replaced with aspartic acid^12^ are perinatal lethal and the survivors display accelerated aging and hypermetabolism suggesting that regulated phosphorylation of DICER1 is critical for normal mammalian development^12^. Interestingly, in a cancer model heterozygous *Dicer1*^*S2D*^ allele together with the heterozygous oncogenic *Kras* G12D (*Kras*^*LA1*^) allele^8^ leads to the generation of LUADs as well as causes the spread of multiple tumors across the animal body^13^.

Canonically, DICER1 functions in the cytoplasm as a RNase enzyme to generate small non-coding RNAs of 21 nucleotides in length, the micro(mi) RNAs^18, 21^. Phosphomimetic DICER1, however, is nuclear in the primary lung tumors in the mouse model with *Kras*^*LA1*^ suggesting that phosphorylation and nuclear localization of DICER1 plays an important role in tumor progression and spread *in vivo*. The discovery that phosphorylated nuclear DICER1 functions with a *Kras* oncogenic mutation to mediate tumor spread suggests that we have uncovered a specific signaling axis of KRAS→ERK→DICER1 cascade that may uniquely drive tumor spread. However, the function of phosphorylated nuclear DICER1 in tumor progression is currently unknown. Here, we investigated the cellular and molecular mechanisms through which phosphorylated nuclear DICER1 affects tumor progression in mice and humans.

We show that DICER1 is phosphorylated and nuclear in human LUADs and correlates with invasion to the lymph nodes. Using genetically modified mouse models, we observe that unlike loss of one copy of *Dicer1*, which regulates tumor onset in a *Kras*^*LA1/+*^ background, phosphomimetic DICER1 only regulates late stages of tumor progression. In addition, and surprisingly, phosphomimetic *Dicer1* regulates late-stage tumor progression independent of epithelial-to-mesenchymal transition (EMT) and miRNA production. Instead, through a combination of single-cell RNA sequencing, *in situ* RNA hybridization and immunofluorescence for validation of single-cell RNA analysis, and ATAC-sequencing, we discovered that the presence of phosphomimetic DICER1 results in a more ‘open’ chromatin state causing reprogramming of the lung tumor alveolar epithelial cells to an endodermal state, such that the tumor cells begin to express gastric gene signatures. Additionally, we observed that grade 3 human LUAD tumors with *KRAS* oncogenic mutations and phosphorylated DICER1 also ectopically express gastric gene signature, suggesting that phosphorylation of DICER1 results in cell reprogramming in mice and humans resulting in progression of LUADs. We propose that the increased tumor cell plasticity due to lineage reprogramming may be the underlying mechanism through which phosphorylated nuclear DICER1 drives late-stage tumor progression.

## Results

### DICER1 is nuclear and phosphorylated in human LUADs

To determine whether DICER1 is phosphorylated in human lung adenocarcinomas (LUADs), we performed immunohistochemistry analysis using anti-phosphorylated DICER1 and anti-phosphorylated ERK antibodies (Methods) on 88 LUAD tumors including 38 (∼43%) samples with *KRAS* oncogenic mutations and 50 (∼57%) without *KRAS* oncogenic mutations. *KRAS* oncogenic mutation status and baseline characteristics of the patient population are presented in Supplementary table 1. Tumors with ≥10% but < 30% of cells with phospho-DICER1 were classified as “low positive”, tumors with ≥30% but <70% of cells as “moderate positive”, ≥70% of cells as “high positive” and <10% of cells with phospho-DICER1 as negative (Methods, Table 1 and Supplementary Fig. 1A). Tumors were simultaneously analyzed for phospho-ERK status following the same parameters as described for phospho-DICER1 (Methods, Table 1 and Supplementary Fig. 1A).

**Table 1.**
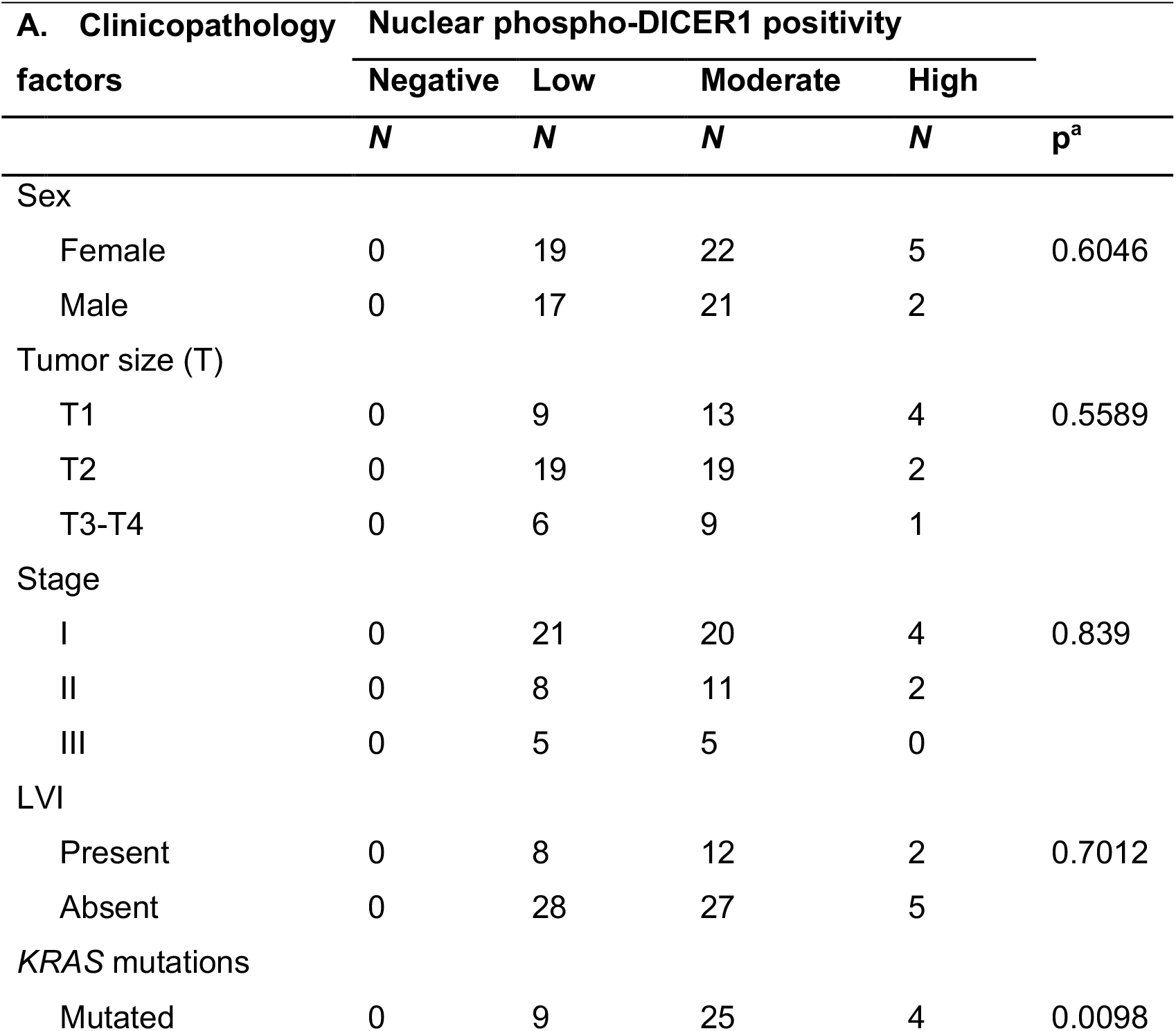
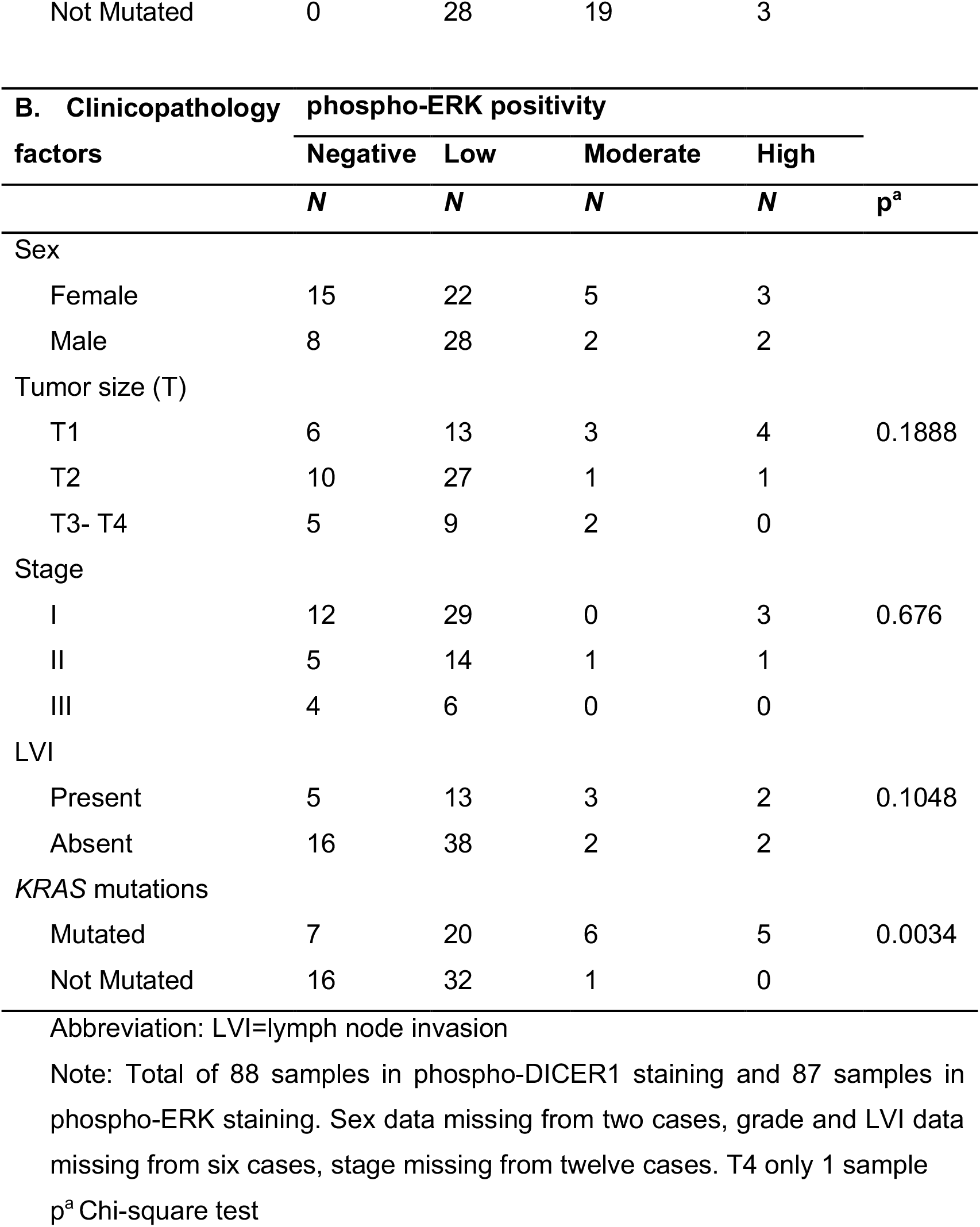
Relationship between phospho-DICER1(A), phospho-ERK (B) and clinicopathological features in LUADs.

| **A. Clinicopathology factors** | **Nuclear phospho-DICER1 positivity** |  |  |  | **p<sup>a</sup>** |
| --- | --- | --- | --- | --- | --- |
|  | **Negative** | **Low** | **Moderate** | **High** |  |
|  | **N** | **N** | **N** | **N** |  |
| Sex |  |  |  |  |  |
| Female | 0 | 19 | 22 | 5 | 0.6046 |
| Male | 0 | 17 | 21 | 2 |  |
| Tumor size (T) |  |  |  |  |  |
| T1 | 0 | 9 | 13 | 4 | 0.5589 |
| T2 | 0 | 19 | 19 | 2 |  |
| T3-T4 | 0 | 6 | 9 | 1 |  |
| Stage |  |  |  |  |  |
| I | 0 | 21 | 20 | 4 | 0.839 |
| II | 0 | 8 | 11 | 2 |  |
| III | 0 | 5 | 5 | 0 |  |
| LVI |  |  |  |  |  |
| Present | 0 | 8 | 12 | 2 | 0.7012 |
| Absent | 0 | 28 | 27 | 5 |  |
| KRAS mutations |  |  |  |  |  |
| Mutated | 0 | 9 | 25 | 4 | 0.0098 |

|  |  |  |  |  |
| --- | --- | --- | --- | --- |
| Not Mutated | 0 | 28 | 19 | 3 |

| B. Clinicopathology factors | phospho-ERK positivity |  |  |  | p<sup>a</sup> |
| --- | --- | --- | --- | --- | --- |
|  | Negative | Low | Moderate | High |  |
|  | N | N | N | N |  |
| Sex |  |  |  |  |  |
| Female | 15 | 22 | 5 | 3 |  |
| Male | 8 | 28 | 2 | 2 |  |
| Tumor size (T) |  |  |  |  |  |
| T1 | 6 | 13 | 3 | 4 | 0.1888 |
| T2 | 10 | 27 | 1 | 1 |  |
| T3- T4 | 5 | 9 | 2 | 0 |  |
| Stage |  |  |  |  |  |
| I | 12 | 29 | 0 | 3 | 0.676 |
| II | 5 | 14 | 1 | 1 |  |
| III | 4 | 6 | 0 | 0 |  |
| LVI |  |  |  |  |  |
| Present | 5 | 13 | 3 | 2 | 0.1048 |
| Absent | 16 | 38 | 2 | 2 |  |
| KRAS mutations |  |  |  |  |  |
| Mutated | 7 | 20 | 6 | 5 | 0.0034 |
| Not Mutated | 16 | 32 | 1 | 0 |  |

Abbreviation: LVI=lymph node invasion
Note: Total of 88 samples in phospho-DICER1 staining and 87 samples in phospho-ERK staining. Sex data missing from two cases, grade and LVI data missing from six cases, stage missing from twelve cases. T4 only 1 sample
p<sup>a</sup> Chi-square test

Phosphorylated DICER1 was positive and nuclear in all LUAD tumors (Fig. 1A-B). Tumors bearing *KRAS* oncogenic mutations correlated with moderate (65% of tumors) to high (10% of tumors) phospho-DICER1 positivity (Fig. 1B) whereas tumors bearing wild type *KRAS* correlated with low (56% of tumors) phospho-DICER1 positivity (Fig 1B). These data demonstrate that positive phospho-DICER1 status correlates significantly with human LUADs bearing *KRAS* mutations.

**Figure 1.**
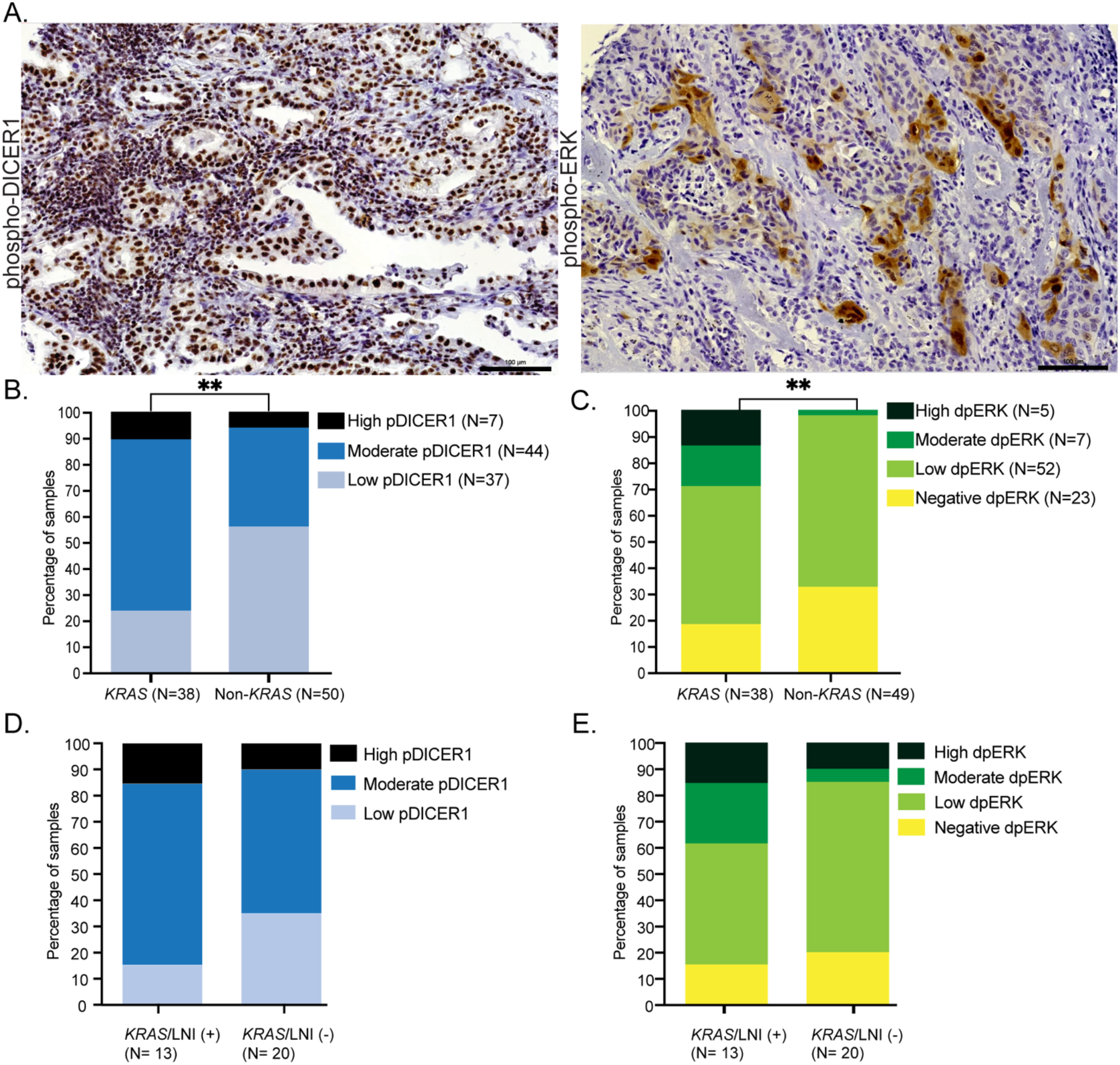
DICER1 is phosphorylated and nuclear in human LUADs. A. Representative images of human LUADs stained with anti phospho-DICER1 (left) and anti phospho-ERK (right) antibodies. 40x, Scale bar =100μm. B. Human LUADs with low, moderate, and high phospho-DICER1 signal plotted as percentages on the y-axis and *KRAS* mutation status on the x-axis. *KRAS*, samples with oncogenic *KRAS* mutations. Non-*KRAS*, samples without mutations in *KRAS*. Samples are classified based on the percentage of cells positive for phospho-DICER1 (pDICER1). Low (<30% positive cells), moderate (≥30% < 70% of positive cells), high (≥70% of positive cells). Chi-square, **p-value= 0.0098. C. Human LUADs with negative, low, moderate, and high phospho-ERK signal plotted by their *KRAS* mutation status. KRAS mutations. Non-*KRAS*, samples without *KRAS* mutations. Samples are classified based on the percentage of cells positive for phospho-ERK (dpERK). Negative (<10% positive cells), low, (≥10% <30% positive cells), moderate (≥30% < 70% of positive cells), high (≥70% of positive cells). Chi-square, **p-value= 0.0034. D. Human LUAD tumors with *KRAS* oncogenic mutations classified based on presence (LNI (+)) or absence (LNI (-)) of lymph node invasion and phospho-DICER1 positive nuclear signal. Chi-square, p-value= 0.2058. E. Human LUAD tumors with *KRAS* oncogenic mutations classified based on presence (LNI (+)) or absence (LNI (-)) of lymph node invasion and phospho-ERK positive nuclear signal. Chi-square, p-value= 0.1048.

Unlike phospho-DICER1 status, only 64 (73%) of the tumors were positive for phospho-ERK (Fig. 1C). Of the tumors with *KRAS* oncogenic mutations that displayed positive phospho-ERK staining, we observed that only 15% of the tumors showed moderate positivity while 13% displayed high level of phospho-ERK signal. The majority of LUAD tumors assayed displayed low phospho-ERK positive signal irrespective of their *KRAS* mutation profile. Surprisingly, however, 18% of the tumors with *KRAS* mutations were negative for phospho-ERK signal. This was surprising because it revealed that phospho-DICER1 and phospho-ERK do not perfectly overlap. This is likely due to a difference in the rate of dephosphorylation between these two markers. However, because positive phospho-ERK is considered a gold standard for aggressive tumors^22^, we asked whether the tumors that were negative for phospho-ERK were invasive.

To determine whether presence or absence of phospho-ERK and phospho-DICER1 correlated with tumor invasion we evaluated tumor pathology and patient clinicopathologic features and observed that phosphorylated DICER1 is more frequently observed in *KRAS* mutated LUADs with invasion to the lymph node (Fig. 1D). A total of 22 LUAD tumors displayed lymph node invasion of which 13 tumors had mutations in *KRAS* and 9 tumors were wild type for *KRAS* (Fig. 1D and Supplementary Fig. 1B). Overall, we observed that a higher proportion of LUAD tumors with lymph node invasion and presence of *KRAS* oncogenic mutations displayed either moderate or high levels of phospho-DICER1 signal. Surprisingly, some tumors with *KRAS* oncogenic mutations and lymph node invasion still showed negative staining for phospho-ERK (Fig. 1E) suggesting that phospho-ERK may be dynamic and a poorer prognostic marker for tumor progression or invasion compared to phospho-DICER1. Collectively, these analyses demonstrate that DICER1 is phosphorylated in human LUADs and correlates with tumor invasion and *KRAS* oncogenic mutations.

### Phosphomimetic-DICER1 causes late-stage tumor progression

*Dicer1* functions as a haploinsufficient tumor suppressor, since loss of one copy of *Dicer1* in a *Kras* oncogenic murine background results in increased tumor size and number as early as 12 weeks in tumorigenesis^16, 17^. Thus, to determine if phosphorylation of DICER1 contributes to tumor initiation or late stage progression in an oncogenic *Kras* background, we utilized the phosphomimetic *Dicer1* murine model (*Dicer1*^*S2D*^) ^12^. In the *Dicer1*^*S2D*^ model, serines at positions 1712 and 1836 are replaced by aspartic acid at the endogenous locus to mimic constitutive phosphorylation. Homozygous *Dicer1*^*S2*D^ mice develop a detrimental aging phenotype resulting in early lethality hindering the study of tumors in these animals^12^. Thus, we crossed double mutants between heterozygous *Dicer1*^*S2*D^ with a heterozygous oncogenic *Kras LA1* murine model^8^ and assayed *Kras*^*LA1/+*^ and *Kras*^*LA1/+*^*;Dicer1*^*S2D/+*^ animals for tumor generation. Animals were evaluated for tumor number, size, and grade at multiple timepoints. *Dicer1*^*S2D/+*^ mice are phenotypically wild type and do not develop tumors as described^12, 13^. *Kras*^*LA1/+*^ mice developed multifocal lung adenomas and adenocarcinomas as described^8^.

At 6 and 12 weeks of age, *Kras*^*LA1/+*^ and *Kras*^*LA1/+*^*;Dicer1*^*S2D/+*^ mice developed numerous hyperplastic lesions and adenomas with no significant difference in the number, size and histological grade (Supplementary Fig.2 A-C), unlike what was observed with loss of one copy of *Dicer1* in a *Kras* oncogenic background^16^. At 24 weeks of age, the *Kras*^*LA1/+*^*;Dicer1*^*S2D/+*^ double mutant mice developed an increase in the number of lung adenocarcinomas with increased cellular atypia and invasion of adjacent bronchial or vessel structures when compared to *Kras*^*LA1/+*^ mice (Fig. 2A and Supplementary Figure 2D). Majority of the adenocarcinomas were organized into papillary like structures with a few solid and lepidic adenocarcinomas. At 34 weeks of age the *Kras*^*LA1/+*^*;Dicer1*^*S2D/+*^ mice displayed tumors across the animal body including lymph node and heart, and thymic lymphomas, in addition to the primary lung adenomas and adenocarcinomas, as previously described^13^ (Fig. 2B-C). *Kras*^*LA1/+*^ single mutant mice displayed no spread of tumors at these timepoints. Altogether, our data indicate that the oncogenic mutation in *Kras* is responsible for tumor initiation and presence of phosphomimetic DICER1 leads to late-stage tumor progression and invasion. These data suggest that the function of DICER1 phosphorylation in mediating tumor progression is distinct from the loss of one copy of *Dicer1*^8^ wherein the latter leads to an enhancement of tumor onset as well as accelerates early stages of tumor development.

**Figure 2.**
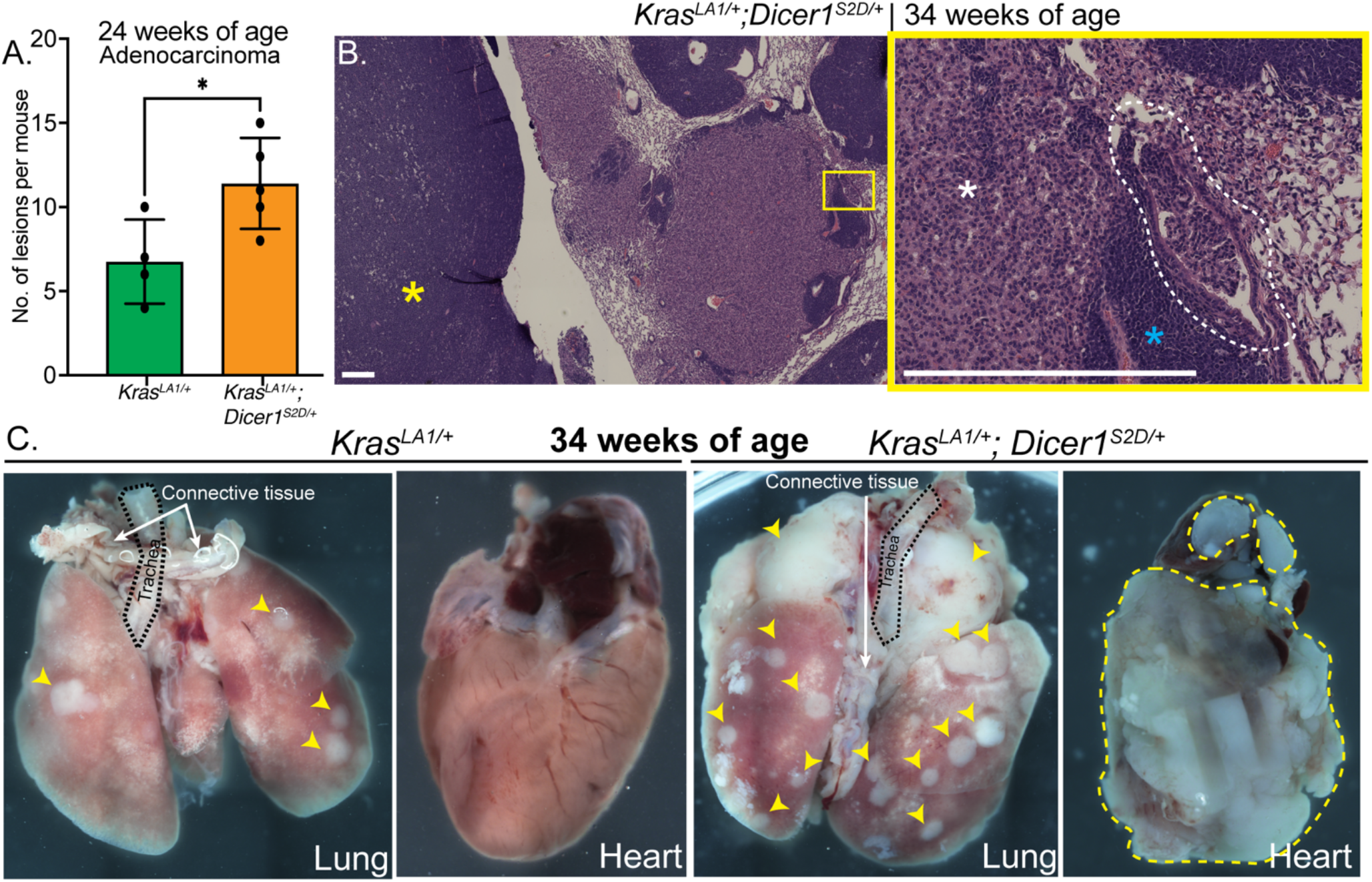
Phosphomimetic *Dicer1* regulates tumor invasion in mice. A. Number of lung adenocarcinoma from *Kras*^*LA1/+*^ (n=4) in green and *Kras*^*LA1/+*^*;Dicer1*^*S2D/+*^ (n=5) in orange at 24 weeks of age. *p-value = 0.03. B. Representative H&E image of the tumor lung adenocarcinomas of *Kras*^*LA1/+*^*;Dicer1*^*S2D/+*^ mice at 34 weeks of age. Left panel shows lower magnification of lung tumors and thymic lymphoma. Right panel shows higher magnification of the invasive lung adenocarcinoma (from left). White asterisk indicates lung adenocarcinoma, yellow asterisk indicates thymic lymphoma and blue asterisk indicates infiltrating lymphocytes. Encircled in white dash-lines are lung adenocarcinoma cells invading nearby bronchial structures. Scale bar = 100μm C. Representative images of gross morphology of a lung and a heart from *Kras*^*LA1/+*^ and *Kras*^*LA1/+*^*;Dicer1*^*S2D/+*^ mice at 34 weeks of age. Yellow arrowheads point to lung tumors. Encircled in yellow is a tumor lesion spreading into and compressing the heart.

Presence of *Kras* oncogenic mutations leads to the formation of lung tumors that originate from alveolar type II (AT2) cells^8^. We observed that *Kras*^*LA1/+*^*;Dicer1*^*S2D/+*^ lung tumors are also AT2 in nature as assessed by the presence of pro-surfactant (pro-SFPC) expression, which marks AT2 cells (Supplementary Fig. 3A). In addition, Aryal. N et al.^13^ showed that the mouse lung tumors display phosphorylated nuclear DICER1 like the human LUADs (Fig. 1A). Thus, together with the genetic analysis we conclude that constitutive phosphorylation of DICER1 downstream of the KRAS oncogenic pathway promotes late-stage tumor progression and lead to formation of invasive tumors *in vivo*.

### Phosphorylated DICER1 does not regulate EMT to mediate tumor invasion

To determine the cellular mechanism through which the presence of phosphomimetic *Dicer1* leads to an increased number of adenocarcinomas and spread of tumors, we evaluated EMT as a mechanism for tumor progression (Methods). Lung tumors from *Kras*^*LA1/+*^*;Dicer1*^*S2D/+*^ and *Kras*^*LA1/+*^ genotypes (1-year-old, n= 5 per genotype) were assayed using immunofluorescence for epithelial and mesenchymal markers (Methods). AT2 cells were stained with the alveolar marker pro-SFPC, which marks mature AT2 cell. We then assayed for Vimentin expression within the AT2 positive tumor cells. We hypothesized that tumor epithelial cells undergoing EMT will display AT2 alveolar markers and co-express the Vimentin mesenchymal marker. However, we observed that AT2 cells did not exhibit Vimentin signal in any of the tumors assayed. Instead, AT2 tumor cells displayed a dual pattern for E-cadherin expression, where some (i) exhibited high levels of E-cadherin staining, consistent with their epithelial nature and some (ii) exhibited low levels of E-cadherin signal (Supplementary Fig. 3A). This variation in E-cadherin staining suggests that the AT2 tumor cell population is heterogeneous with regards to E-cadherin expression. Yet, even in AT2 tumor cells with low E-cadherin expression, we did not observe any Vimentin expression. In addition, we observed that the lung tumors from *Kras*^*LA1/+*^*;Dicer1*^*S2D/+*^ and *Kras*^*LA1/+*^ mice displayed similar percentages of Vimentin positive cells irrespective of their E-cadherin status (Supplementary Fig. 3B). These data demonstrate that the tumor cells are not changing from epithelial (E-cadherin) to mesenchymal (Vimentin positive) state and that the tumor cells bearing E-cadherin are mutually exclusive from cells that displayed Vimentin expression. In general, cells that expressed Vimentin were likely immune and fibroblast cells. Thus, given the lack of Vimentin staining in AT2 cells and no difference in the number of Vimentin positive cells between both the genotypes we conclude that phosphomimetic DICER1 induced tumor progression is independent of EMT.

### Phosphorylated DICER1 results in expression of gastrointestinal genes in lung tumors

During the EMT analysis, we observed a heterogeneity in E-cadherin expression within the AT2 tumor cell populations. Thus, we hypothesized that phosphomimetic DICER1 likely generates sub-clonal populations of cells that contribute to tumor progression or invasion. To determine whether phosphomimetic DICER1 leads to the generation of sub-clonal populations of tumor cells that contribute to tumor invasion in the *Kras*^*LA1/+*^*;Dicer1*^*S2D/+*^ relative to *Kras*^*LA1/+*^ animals, we performed single-cell RNA sequencing (scRNA-Seq) on the lung tumors from both the genotypes at 39 weeks of age (Methods). We isolated and sequenced 27 lung tumors (20,214 cells) from *Kras*^*LA1/+*^ mice and 23 lung tumors (18,754 cells) from *Kras*^*LA1/+*^*;Dicer1*^*S2D/+*^ mice. In addition, we sequenced lung tissue (7,288 cells) from *Dicer1*^*S2D/+*^ mouse to validate the expression of canonical lung markers. In the lung tumors, we identified a total of 18,816 cells clustered into three major cell populations belonging to the epithelial, immune, and mesenchymal lineages (which includes endothelial cells) (Supplementary Fig. 4A-B). Cells positive for *Epcam, Cdh1, Sfta2, Lamp3, Scl34a2, Rtkn2, Pdpn, Hopx, Ager, Scgb1a1, Scgb3a2* and *Foxj1* were classified as epithelial tumor cell populations and analyzed by sub-clustering according to the markers that each cell expressed (Methods, Supplementary Fig. 4C). Epithelial cell populations included alveolar type I (AT1), AT2, Clara and Ciliated cells (Supplementary Fig. 4C). Within the AT2 cell population, we observed at least four sub-populations, (i) those positive for *Cdh1* (E-cadherin) and (ii) those negative for *Cdh1* (3-fold-change difference) (Supplementary Fig. 5A), consistent with the immunofluorescence analysis (above). We named these two populations as AT2.a and AT2.b respectively (Figure 3A-B and Figure 3D, Supplementary Fig. 5A and 5E). (iii) the third sub-cluster of epithelial cells expressed AT2 cell markers but, surprisingly, also expressed gastrointestinal genes including *Gkn2, Ly6d, Apoe, A2ml1, Emp1* (Fig. 3A-D and Supplementary Fig. 5B-D). Expression of the gastrointestinal genes is unexpected because gastric genes are normally not expressed in the lung. Because these tumor cells express signatures of endodermal genes, AT2 cells and *Cdh1*, we named this sub-cluster as Alveolar_ Endodermal cells. (iv) the final sub-cluster of tumor epithelial cells displayed low expression of AT2 cell markers and no expression of *Cdh1* or *Lamp3* but still expressed endodermal genes. This sub-cluster was named ‘Endodermal cells’ (Fig. 3A-D and Supplementary Fig. 5B-D).

**Figure 3.**
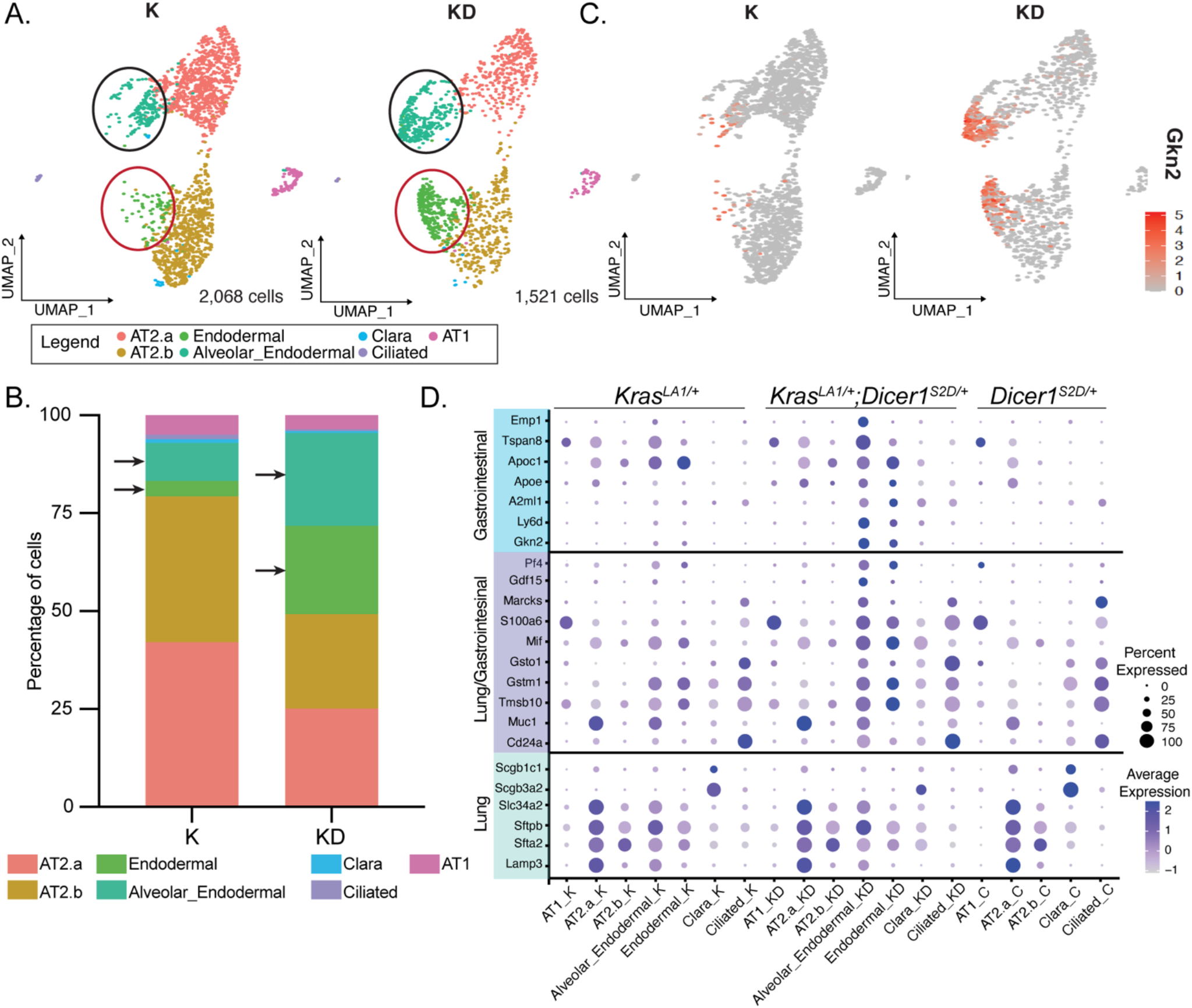
Enrichment of gastrointestinal genes in lung tumor cells with phosphomimetic *Dicer1*. A. UMAP graphs of the clusters identified in the lung tumor epithelial cells from single-cell RNA sequencing analysis of *Kras*^*LA1/+*^ (K, n=2) and *Kras*^*LA1/+*^*;Dicer1*^*S2D/+*^ (KD, n=2) mice. Each dot represents a cell. Color codes describe the various cell types as described in the legend. Green and dark green cell clusters (marked by the black and red circles) are tumor cells with expression of gastrointestinal genes. B. Percentages of the distinct cell types identified in *Kras*^*LA1/+*^ (K, n=2) and *Kras*^*LA1/+*^*;Dicer1*^*S2D/+*^ (KD, n=2) lung tumors. Arrows pointing to Alveolar_Endodermal and Endodermal clusters. C. UMAP showing the relative expression for Gastrokine-2 (*Gkn2*) gene in red within the tumor epithelial cell cluster from *Kras*^*LA1/+*^ (K, n=2) and *Kras*^*LA1/+*^*;Dicer1*^*S2D/+*^ (KD, n=2). D. Dot plot showing gene expression of lung and gastrointestinal gene signatures. The size of the dots corresponds to the percentage of cells (on the x-axis) that express the genes displayed on the y-axis. The color density corresponds to the fold expression of each gene on the y-axis. K labels lung tumor cells from *Kras*^*LA1/+*^, KD labels lung tumor cells from *Kras*^*LA1/+*^*;Dicer1*^*S2D/+*^ and C labels the control lung cells from *Dicer1*^*S2D/+*^.

Together, these data reveal that the lung tumors from *Kras*^*LA1/+*^*;Dicer1*^*S2D/+*^ mice bear tumor epithelial cell populations comprised of 25% of AT2.a cells, 24% of AT2.b cells, 23% Alveolar_ Endodermal and 22% Endodermal clusters cells. In contrast, in the *Kras*^*LA1/+*^ single mutant lung tumors, over 42% and 37% of the tumor epithelial cells are comprised of AT2.a and AT2.b tumor cells respectively, whereas Alveolar_ Endodermal and Endodermal clusters represent only 9.7% and 3.9%, respectively (Fig. 3B). The significant enrichment in the Alveolar_ Endodermal and Endodermal clusters in the *Kras*^*LA1/+*^*;Dicer1*^*S2D/+*^ relative to the *Kras*^*LA1/+*^ tumors suggests that phosphorylated nuclear DICER1 in the lung tumors leads to ectopic expression of gastric and other endodermal genes which creates sub-clonal populations of tumor cells that alter tumor cell plasticity and likely drive invasion.

### Phosphorylated DICER1 in murine lung tumors alters alveolar type II identity

To validate the expression of gastric genes identified from the scRNA-sequencing analysis (above) we performed Hairpin Chain Reaction (HCR)-RNA FISH (Molecular instruments, Methods) and immunofluorescence staining on lung tumors of *Kras*^*LA1/+*^ and *Kras*^*LA1/+*^*;Dicer1*^*S2D/+*^ mice (Methods). We tested for gastrokine-2 (*Gkn2*) expression because (i) it is upregulated in the *Kras*^*LA1/+*^*;Dicer1*^*S2D/+*^ lung tumor cells, (ii) *Gkn2* is expressed in the wild type stomach but not lungs and (iii) a subset of poorly characterized human lung cancers known as mucinous adenocarcinoma display GKN1 expression^23^. Although, the mechanism for the latter remains unknown. Consistent with the scRNA-sequencing analysis, we observed the presence of *Gkn2* mRNA and protein in lung tumor epithelial cells from *Kras*^*LA1/+*^*;Dicer1*^*S2D/+*^ animals (Fig. 4A-C and Supplementary Fig. 6). Furthermore, the number of lung tumors with populations of epithelial cells that express *Gkn2* was significantly higher in the *Kras*^*LA1/+*^*;Dicer1*^*S2D/+*^ relative to the *Kras*^*LA1/+*^ mice (Fig. 4B). We also identified sub-populations of tumor cells which expressed *Gkn2* but did not express pro-surfactant, a marker for mature AT2 cells (Fig. 4A and 4C, white head arrows)^23, 24^. These data suggest that this sub-clonal population of tumor epithelial cells loses their AT2 mature marker and thus AT2 cell identity while gaining the expression of gastric genes. Finally, we observed that *Kras*^*LA1/+*^*;Dicer1*^*S2D/+*^ tumors with *Gkn2* expression were poorly differentiated with features of desmoplasia, secretion of mucus, disorganized cells with variable nuclear size and shape, and invasion to bronchial structures when compared to tumors that are *Gkn2* negative, which are differentiated and organized into either papillary, solid or lepidic types (Fig. 4C). Thus, we conclude that phosphomimetic DICER1 in the lung tumors alters AT2 cell identity and causes onset of gastric like cell fate together leading to a poorly differentiated tumor in the *Kras*^*LA1/+*^*;Dicer1*^*S2D/+*^ mice.

**Figure 4.**
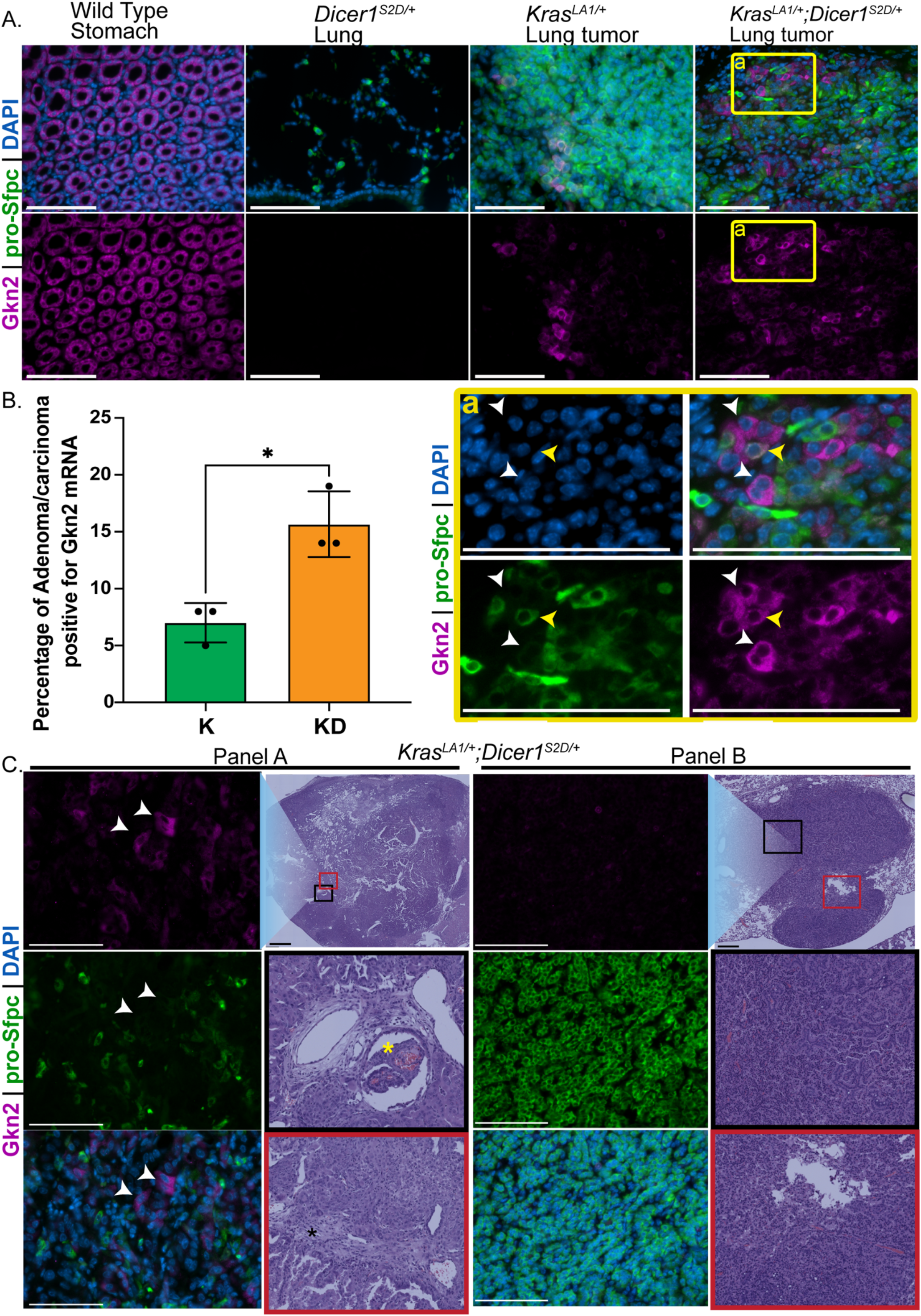
Phosphomimetic *Dicer1* alters the identity of AT2 tumor cells and enhances tumor cell plasticity. A. Representative images of *Gkn2* RNA (in magenta), pro-*Sfpc* RNA (in green) and nuclei (DAPI in blue) in wild type stomach (positive control for *Gkn2*), *Dicer1*^*S2D/+*^ lung (positive control for pro-*Sfpc*) and lung tumors from *Kras*^*LA1/+*^ and *Kras*^*LA1/+*^*;Dicer1*^*S2D/+*^ mice. Panel (a) shows zoom in of the tumor region indicated by the yellow squares of *Kras*^*LA1/+*^*;Dicer1*^*S2D/+*^ animals. Yellow arrowhead points to cells that co-express *Gkn2* and pro-*Sfpc*. White arrowhead points to cells that only express *Gkn2*. Scale bar = 50μm B. Percentage of lung tumors with epithelial cell populations positive for *Gkn2* mRNA in *Kras*^*LA1/+*^ (K, N=3 mice, n= 45 lung tumors) and *Kras*^*LA1/+*^*;Dicer1*^*S2D/+*^ (KD, N=3 mice, n= 44 lung tumors) mice. C. Correlation between *Gkn2* mRNA staining and the tumor histology of *Kras*^*LA1/+*^*;Dicer1*^*S2D/+*^ lung tumors (n= 3). Each panel shows *Gkn2* mRNA in magenta, pro-*Sfpc* in green and DAPI in blue in left column and H&E staining in right column. Panel A shows cells that don’t express pro-*Sfpc* while expressing *Gkn2* (mark by white arrowhead) in a lung adenocarcinoma that displayed invasion to the bronchial structures (yellow asterisk), desmoplasia (black asterisk) and poorly organized tumor cells. Panel B shows the lack of *Gkn2* expression in tumor cells that express pro-*Sfpc* in a localized and well differentiated lung adenocarcinoma. White scale bar = 50μm and black scale bar = 100μm

### Phosphorylated DICER1 correlates with expression of gastrokine-2 in human LUADs

Next, we assessed whether human LUADs with oncogenic *KRAS* mutations and phospho-DICER1 expression also expressed *GKN2* RNA. Thus, we performed HCR-RNA FISH analysis on the human tumor tissue microarrays assayed above. We observed the presence of *GKN2* positive tumor cells in poorly differentiated grade 3 stage IIB lung adenocarcinoma (Fig. 5); the tumor also displayed 50% (moderate) positive signal for phospho-DICER1. Notably, tumor cells expressing *GKN2* were positive for nuclear phospho-DICER1 (Fig. 5). These data suggest that phosphorylated nuclear DICER1 in the lung tumors correlates with changes in gene expression that in turn results in cellular reprograming and increased tumor plasticity. Collectively, the mouse and human data suggest that phosphorylation of DICER1 in lung tumors bearing oncogenic *KRAS* mutations is altering tumor cell identity leading to increased lineage and cellular plasticity as the mechanism for tumor progression.

**Figure 5.**
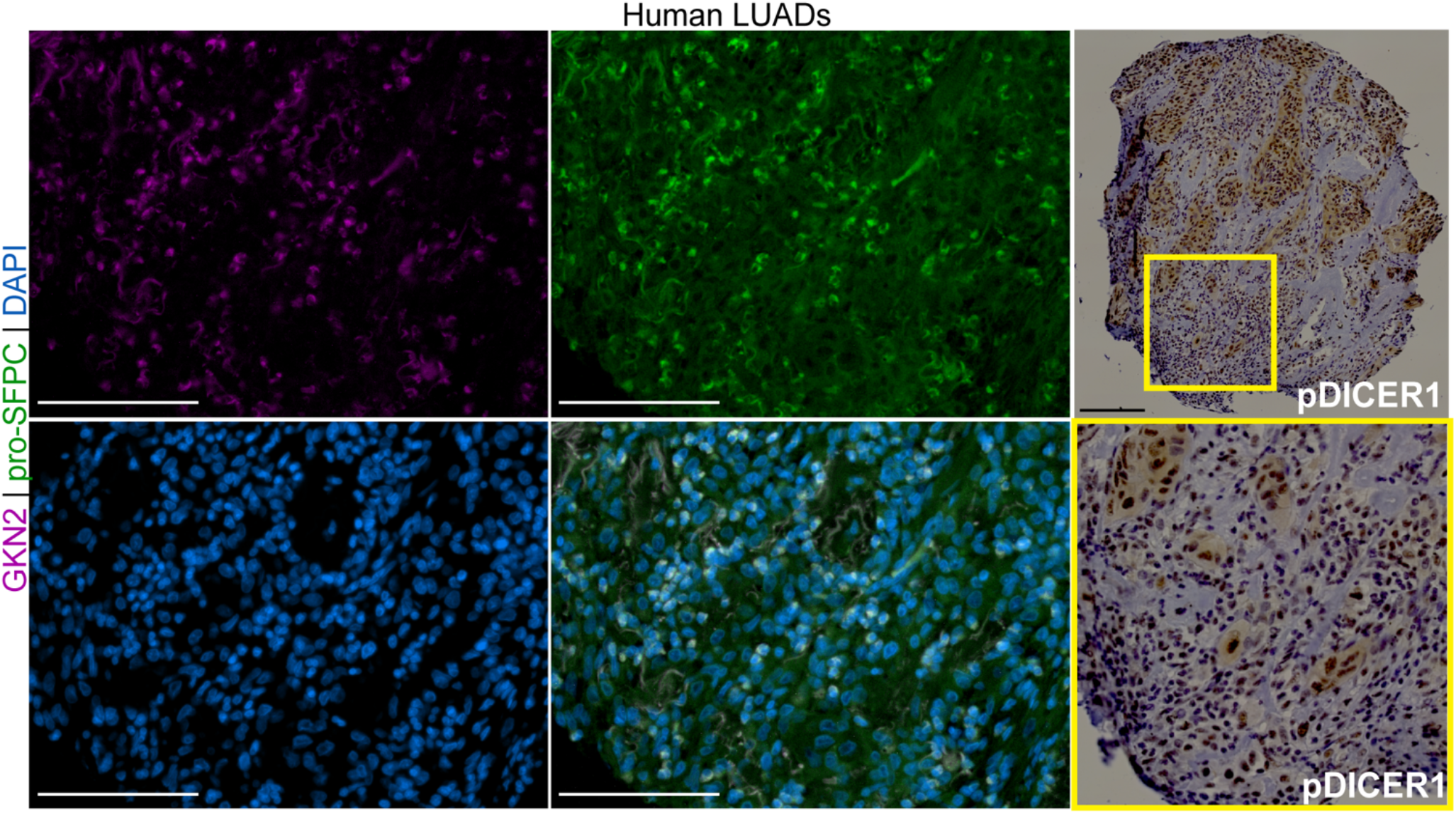
Phosphorylated nuclear DICER1 correlates with expression of GKN2 in human LUADs. Representative images of human LUAD classified as grade 3, stage IIB with positive signal for *GKN2* mRNA. Right panel: human LUAD with positive signal for phospho-DICER1 antibody (in brown). White scale bar = 50μm; Black scale bar= 100μm

### Phosphorylated nuclear DICER1 does not regulate miRNA production to control gene expression

DICER1 is a RNase enzyme that generates mature miRNAs for post-transcriptional regulation of genes^18^. To determine if phosphorylation of DICER1 in lung tumors affects the production of miRNAs and consequently mRNA expression, we assayed for mature miRNAs in the lung tumors from *Kras*^*LA1/+*^ (n= 4) and *Kras*^*LA1/+*^*;Dicer1*^*S2D/+*^ (n= 5) mice using the Nanostring nCounter mouse miRNA expression array (Methods). Overall, we observed that 97.5% of the miRNAs were unchanged in the lung tumors between the two genotypes (n= 577 miRNAs) (Supplementary Fig. 7A). To determine whether the remaining 2.5% of miRNAs that showed either upregulation or downregulation in the *Kras*^*LA1/+*^*;Dicer1*^*S2D/+*^ mice (cut off FC ≥ 2 or ≤ -1.5; none showed ≤ - 2) were known oncogenic or tumor suppressor miRNAs, we zoomed in on each individual miRNA (Supplementary Fig. 7B). We found that miR-434-3p, miR-148b, miR-15b, miR-322 and miR-150 miRNAs were significantly upregulated in the *Kras*^*LA1/+*^*;Dicer1*^*S2D/+*^ when compared to the *Kras*^*LA1/+*^ mice (Supplementary Fig. 7B), while miR-148a, miR-1937, miR-30a and miR-429 were significantly downregulated in the *Kras*^*LA1/+*^*;Dicer1*^*S2D/+*^ animals when compared to the *Kras*^*LA1/+*^ mice (Supplementary Fig. 7B). However, we did not identify any common patterns in their function from literature, or seed sequences, which target mRNAs for regulation (Supplementary Fig. 7C). For example, in our data set miR-148b, miR-15b and miR-150 are upregulated in the *Kras*^*LA1/+*^*;Dicer1*^*S2D/+*^ tumor when compared to the *Kras*^*LA1/+*^ tumor, yet, in literature while miR-15b and miR-150 are thought to be upregulated in lung cancer cell lines miR-148b is downregulated^25-30^. Similarly, miR-148a and miR-30a and miR-429 were downregulated in *Kras*^*LA1/+*^*;Dicer1*^*S2D/+*^ when compared to *Kras*^*LA1/+*^ tumors yet, in literature miR-429 upregulation was observed in lung cancer cell lines^31-34^. Altogether given the minimal change in miRNA expression between the two genotypes, lack of any seed sequence conservation and lack of correlation between these miRNAs and cancer progression, we suggest that phosphomimetic DICER1 does not significantly impact miRNA production in lung tumors.

### Phosphorylated nuclear DICER1 regulates the chromatin state to control gene expression and tumor progression

To determine the mechanism through which phosphorylated nuclear DICER1 may regulate gene expression (as assessed by scRNA-seq analysis) independent of miRNA production, we performed a chromatin immunoprecipitation analysis with total DICER1 antibody on lung tissue from *Dicer1*^*S2D/S2D*^ (Methods) because phosphorylated DICER1 resides in the nucleus. We observed that nuclear DICER1 does not directly bind to the chromatin (Supplementary Fig. 8A). In the literature, however, nuclear DICER1 has been implicated to function with chromatin remodeling factors to control chromosome segregation^35^. And since nuclear phosphorylated DICER1 did not directly bind to the chromatin we next hypothesized that it may regulate gene expression through binding indirectly to the chromatin. Thus, we performed a global ATAC-sequencing analysis on lung tumors from *Kras*^*LA1/+*^*;Dicer1*^*S2D/+*^ and *Kras*^*LA1/+*^ animals (Methods) to assess whether presence of phosphomimetic DICER1 leads to a change in chromatin accessibility or compaction which may in turn lead to the altered gene expression observed (scRNA-seq analysis above) in the lung tumors.

We identified 43,137 and 42,524 peaks, on average, corresponding to open chromatin in the lung tumors from *Kras*^*LA1/+*^ and *Kras*^*LA1/+*^*;Dicer1*^*S2D/+*^ mice, respectively. While the peak distribution seems similar between the two genotypes (Supplementary Fig. 8B-D), we observed that the presence of phosphomimetic nuclear DICER1 in the lung tumors results in changes in chromatin compaction in the *Kras*^*LA1/+*^*;Dicer1*^*S2D/+*^ tumors relative to the *Kras*^*LA1/+*^ tumors at several loci throughout the genome (Fig. 6A-E and Supplementary Fig. 8E). Specifically, we observed that the lung tumors from the *Kras*^*LA1/+*^*;Dicer1*^*S2D/+*^ mice display peaks corresponding to open chromatin at the beginning of the *Gkn2* genomic locus which are absent from the *Kras*^*LA1/+*^ lung tumors as well as the wild type lung (Fig. 6A). Similarly, we observed peaks corresponding to open chromatin in the lung tumors of *Kras*^*LA1/+*^*;Dicer1*^*S2D/+*^ mice at other gastrointestinal genes such as *A2ml1, Hnf4a* and *Ctse* locus; which were also absent in the *Kras*^*LA1/+*^ lung tumors and wild type lung (Fig. 6B-D). The state of the open chromatin at loci from gastrointestinal genes in the tumors bearing phosphomimetic DICER1 supports the observed increase in gene expression of the gastrointestinal genes in *Kras*^*LA1/+*^*;Dicer1*^*S2D/+*^ lung tumors (Fig. 3-4, Supplementary Fig. 4-6).

**Figure 6.**
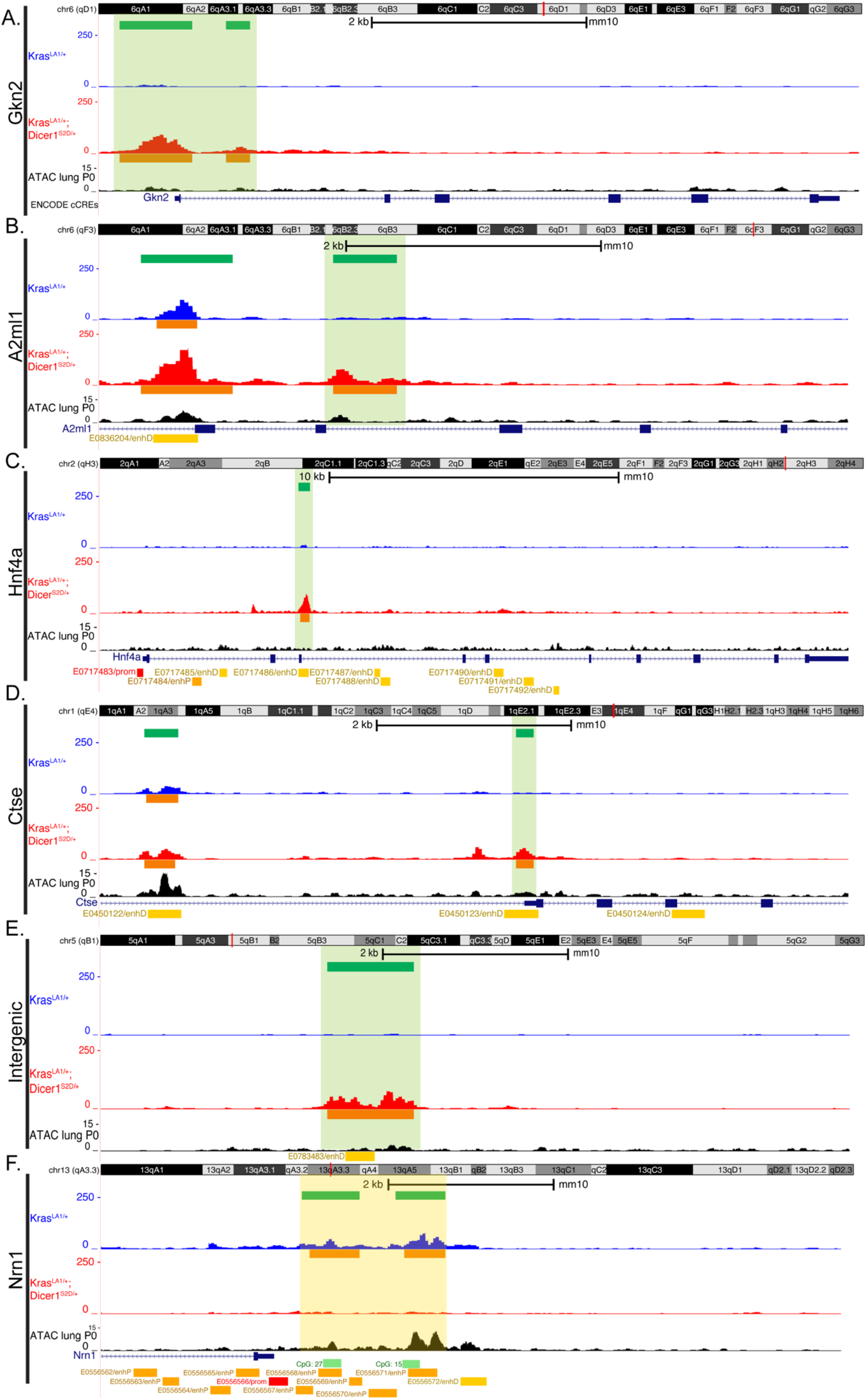
Phosphomimetic nuclear DICER1 affects chromatin compaction in the lung tumors. A. Representative histogram showing ATAC-sequencing results for *Kras*^*LA1/+*^ (in blue), *Kras*^*LA1/+*^; *Dicer1*^*S2D/+*^ (in red) lung tumors, and wild type mouse lung (in black) at the *Gkn2* genomic locus. B. Representative histogram showing ATAC-sequencing results for *Kras*^*LA1/+*^ (in blue), *Kras*^*LA1/+*^; *Dicer1*^*S2D/+*^ (in red) lung tumors, and wild type mouse lung (in black) at the *A2ml1* genomic locus. C. Representative histogram showing ATAC-sequencing results for *Kras*^*LA1/+*^ (in blue), *Kras*^*LA1/+*^; *Dicer1*^*S2D/+*^ (in red) lung tumors, and wild type mouse lung (in black) at the *Hnf4a* genomic locus. D. Representative histogram showing ATAC-sequencing results for *Kras*^*LA1/+*^ (in blue), *Kras*^*LA1/+*^; *Dicer1*^*S2D/+*^ (in red) lung tumors, and wild type mouse lung (in black) at the *Ctse* genomic locus E. Representative histogram showing ATAC-sequencing results for *Kras*^*LA1/+*^ (in blue), *Kras*^*LA1/+*^; *Dicer1*^*S2D/+*^ (in red) lung tumors, and wild type mouse lung (in black) an intergenic genomic region in chromosome 5. For each panel: ATAC-sequencing from wild type mouse lung was retrieved from ENCODE Consortium 3 database. Green bars show the “Merged Regions” as described in Methods. Orange bars shows individual Intervals for each sample as described in Methods. For each gene, exons are represented as rectangular bars and introns as lines with arrows (pointing the direction of transcription). Regulatory elements in the mouse genome as identified by the ENCODE Registry of candidate cis-Regulatory Elements (cCREs) database are shown in the bottom of each panel. Highlighted in light green are regions where peaks were present in the *Kras*^*LA1/+*^*;Dicer1*^*S2D/+*^ lung tumors and absent in the *Kras*^*LA1/+*^ lung tumors. Highlighted in light yellow are regions absent in the *Kras*^*LA1/+*^*;Dicer1*^*S2D/+*^ lung tumors and present in the *Kras*^*LA1/+*^ lung tumors.

In addition to specific gastrointestinal loci with open chromatin, we also observed peaks at *Hnf4a, Ctse* and several intergenic regions with signatures of distal-enhancers (Fig. 6B-E). These data suggest that the presence of phosphomimetic nuclear DICER1 induces open chromatin state in regions with regulatory elements such as enhancers, which in turn leads to changes in gene expression. Finally, albeit less frequently, we observed peaks in the *Kras*^*LA1/+*^ lung tumors that were low or absent in the *Kras*^*LA1/+*^*;Dicer1*^*S2D/+*^ lung tumors, such as in *Nrn1* and *Fezf1* locus (Fig. 6F and Supplementary Fig. 8E). Absence of peaks or low peak signal in *Kras*^*LA1/+*^*;Dicer1*^*S2D/+*^ lung tumors relative to *Kras*^*LA1/+*^ animals suggests a closed chromatin state in the presence of phosphomimetic nuclear DICER1. Further analysis of the regions that displayed closed chromatin in the *Kras*^*LA1/+*^*;Dicer1*^*S2D/+*^ revealed that these genomic regions bear CpG islands. These data suggest that these regions may be obscured from DNA methylation in the closed conformation further causing a change in gene expression. Altogether, these data demonstrate that the presence of phosphomimetic DICER1 in the lung tumors bearing *Kras* oncogenic mutation leads to alteration in chromatin compaction in regions with regulatory elements, which in turn leads to the ectopic expression of specific gastrointestinal genes in the tumor epithelial cells resulting in a tumor with increased lineage plasticity as a mechanism for tumor progression.

## Discussion

Our work reports on the role of DICER1 phosphorylation in mediating tumor progression and spread, in a manner that is distinct from DICER1’s function as a haploinsufficient tumor suppressor. Below, we discuss the role of phosphorylated nuclear DICER1 in regulating cellular reprogramming and lineage plasticity as a mechanism for cancer progression, how phosphorylated nuclear DICER1 functions independently of miRNA production, and the potential use of phosphorylated DICER1 antibody as a prognostic marker for detection of malignant human cancers.

### Phosphorylated nuclear DICER1 reprograms tumor cell identity and leads to tumor progression and spread

In its canonical function, DICER1 functions in the cell cytoplasm to process small non-coding RNAs (the miRNAs and siRNAs) and regulate gene expression post-transcriptionally^18^. Loss of one copy of DICER1, in this context, was shown to cooperate with the *Kras* oncogenic mutation G12D and lead to tumor onset and initiation^16, 17^. The mechanism through which DICER1 functions to control tumor onset was proposed to be through the regulation of miRNA production^16^. As a haploinsufficient tumor suppressor, DICER1 has also been shown to function as a regulator of metastasis through the regulation of miRNAs^36^. In these situations, the miRNAs produced lead to EMT as a mechanism for tumor spread. We uncovered a unique role for post-translationally modified phosphorylated DICER1 in regulating only late-stage tumor progression in a manner independent of EMT. We find that phosphorylated nuclear DICER1 causes reprogramming of the lung tumor epithelial alveolar cells to an endodermal state leading to the expression of gastric genes. We propose a model wherein oncogenic mutations in *Kras* boosts alveolar proliferation and initiates the tumor, then phosphorylation of DICER1 through the KRAS-ERK signaling axis results in tumor progression and spread by altering the identity of alveolar type 2 cells (Fig. 7). We show that phosphorylated nuclear DICER1 causes the AT2 cells to assume either an “intermediate stage” where the tumor cells express signatures for both alveolar and gastrointestinal genes or the tumor cells are fully altered in their identity and lose the mature alveolar markers. Seemingly, the ability of phosphorylated nuclear DICER1 to reprogram alveolar cells may be a consequence of hijacking or de-repressing the differentiation program which is laid down during embryonic lung development; lungs originate from the embryonic foregut during endoderm formation, which also gives rise to the esophagus, stomach, liver, and the pancreas^24, 37^. In addition, both *Dicer1* and *Kras* are essential for normal lung development in mice^38^. Loss of *Dicer1* results in defective branching morphogenesis and alveolar differentiation^38^, and KRAS promotes lung branching^39-42^. Thus together, these observations suggest that KRAS-ERK-DICER1 signaling axis may play a central role during normal lung development and it is likely that this role is dysregulated during tumorigenesis.

**Figure 7:**
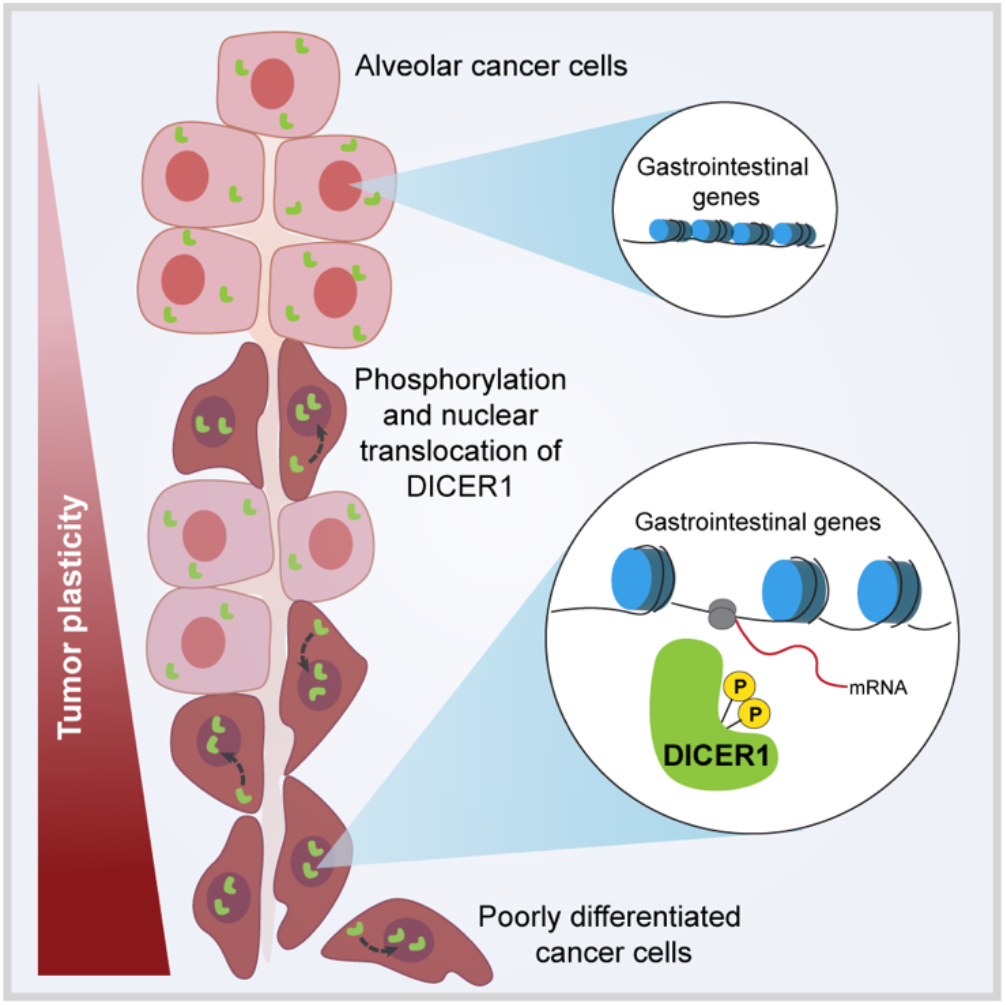
Phosphorylated nuclear DICER1 regulates late-stage tumor progression through reprogramming the chromatin and leading to increased cell plasticity.

Notably, *Nkx2-1*, a transcription factor important for alveolar differentiation when deleted from an oncogenic *Kras* lung adenocarcinoma in a genetically modified mouse model also results in ectopic expression of gastrointestinal genes (including gastrokine-1) and downregulation of alveolar markers such as surfactant (pro-*Sfpc*)^23, 43, 44^. Similarly, coupling loss of *Nkx2-1* with overexpression of transcription factor *Sox2* in murine models promoted the formation of squamous lung cancer with features of esophageal differentiation^43^. Thus, suggesting that a change in cell fate is a common mechanism in a subset of lung cancer. Interestingly, however, loss of *Nkx2-1* in a *Kras* wild type lung failed to express gastrokine-1 suggesting that KRAS may activate specific signaling pathway that augment changes to the differentiation state, in a mechanism independent of, or redundant with, *Nkx2-1* function. We propose that phosphorylation of DICER1 may function in parallel with programs such as *Nkx2*.*1* to regulate alveolar cell identity and result in lung tumors with enhanced plasticity as a mechanism for late-stage tumor progression and invasion. Interestingly, prior to our study alteration in alveolar cell identity to a gastric cell like lineage were uniquely observed with the loss or gain of transcription factors that govern embryonic cell-fate specification. We report phosphorylation of DICER1 which is a RNAse enzyme as a novel regulator in this process which functions to promote changes in cell-lineage through a unique mechanism such as altering the chromatin compaction on a global level.

### Phosphorylated nuclear DICER1 alters chromatin compaction to mediate lineage plasticity of tumor cells aiding their progression

Unlike its canonical function in the cytoplasm, wherein DICER1 regulates miRNA production, to drive tumor initiation and metastasis in cancers^16, 36^, we show that phosphomimetic DICER1 drives tumor progression in a mechanism independent of miRNA function (Supplementary Fig. 7). Instead, we find that the presence of phosphomimetic DICER1 in the lung tumors with *Kras* oncogenic mutations correlates with global chromatin changes (Fig. 6 and Supplementary Fig. 8). Interestingly, these changes lead to opening and altering of chromatin compaction in areas overlapping with regulatory elements in the genome. In support of this, we identified regions around gastrokine-2 that are ‘open’ in the presence of phosphomimetic DICER1 relative to wild type lungs (Fig. 6), suggesting that an aberrant gene expression program is driven in this instance by changes to the chromatin landscape. We find that DICER1 does not directly bind to the chromatin (Supplementary Fig. 8A), thus it is likely that in the nucleus phosphorylated DICER1 partners with novel interactors that affect chromatin remodeling. Our data lead to the model that once phosphorylated, DICER1 translocates to the nucleus where it affects chromatin compaction, which in turn leads to the expression of gastrointestinal genes in alveolar tumor cells and changes to cellular identity and enhanced tumor plasticity (Fig. 7).

In support of these observations, DICER1 has been shown to function with the bromodomain transcription co-activator BRD4 to modulate genome stability and transcription in mouse embryonic stem cells (ES cells)^35^. Gutbrod et al., show that loss of DICER1 in mouse ES cells leads to defects in proliferation and alterations in transcription of centromeric satellite repeats; defects that can be rescued by loss of BRD4 or inhibiting RNA polymerase II^35^. In yeast, DICER1 have been shown to act on RNA substrates to release RNA polymerase II from sites where the transcription and replication machinery may collate^45^. These studies support a role for DICER1 in chromatin reprograming. Yet, the exact molecular mechanism through which nuclear DICER1 modulates chromatin compaction and transcription remains unknown. Because we find that phosphomimetic nuclear DICER1 does not directly bind to the DNA to regulate chromatin compaction (Supplementary Fig. 8A), we speculate that phosphorylated DICER1 is likely recruited to the genome through either DNA:RNA hybrids or RNA intermediary, as shown for nuclear DICER1 in Hek293 cells^46, 47^, or through intermediary protein partners.

Taken together, because DICER1’s function as a haploinsufficient tumor suppressor occurs through loss of miRNA production^16^ and phosphorylation of DICER1 promotes late-stage of tumor progression in a mechanism that is independent of the miRNAs, our study demonstrates that phosphorylation is modulating or activating non-canonical functions of DICER1, such as chromatin compaction. These data provide new directions to the field to help identify the molecular partners through which phosphorylation of DICER1 modulates chromatin compaction in cancers.

### Phosphorylated nuclear DICER1 as a prognostic marker for aggressive lung adenocarcinomas

Cancer development can be divided into two main phases, an initiating phase driven by mutations or genomic alterations that start the tumor formation and a progression phase driven and sustained by further genomic, metabolic or epigenetic changes that drive metastasis^48, 49^. 20-30% of lung adenocarcinomas are initiated due to alveolar cells acquiring somatic oncogenic mutations in *KRAS*^2-5^. As the tumor progresses, cells accumulate additional epigenetic and molecular alterations that modulate cell behavior and, in some cases, cellular identity. While early detection of mutations, which lead to tumor onset are important to screen for cancer prevention, not all mutations lead to cancer progression, and often appear seemingly dormant for years before causing obvious drastic effects. Thus, it is critical to identify the genetic or epigenetic changes that lead to tumor progression and metastasis early enough such that patients can be stratified for effective therapeutic options. Here, we show that phosphorylated nuclear DICER1 correlates with aggressive human lung adenocarcinomas bearing *KRAS* oncogenic mutations. Experimentally, in the mouse models, we demonstrate that phosphorylated nuclear DICER1 drives late-stage tumor progression by affecting alveolar identity; a phenomenon we then observe in human lung adenocarcinomas as well. Most importantly, we show that phosphorylated DICER1 is more frequently detected by immunohistochemistry than active ERK in human lung adenocarcinomas; and in some tumors phosphorylated ERK was absent yet phosphorylated DICER1 was present, and the tumors were aggressive and invasive in nature. Collectively, these data suggest that phosphorylated nuclear DICER1 may represent a potential biomarker for early detection of metastatic cancers and opens the field to develop ways to block DICER1 phosphorylation and nuclear translocation as a means for therapy.

## Author contributions

R.R.C and S.A designed the research, analyzed the data, and wrote the manuscript. R.R.C, S.C and J.S performed key research experiments and contributed figures. All authors read and approved the paper.

## Acknowledgements

We thank Dr. Guillermina Lozano for providing the mice, expertise in analysis and for critical comments on the manuscript. We thank Dr. Nicholas Navin and Dr. Jichao Chen for help with single-cell RNA sequencing protocols and analysis. We thank Dr. Elizabeth Whitley for performing mouse pathology examination and diagnosis. We thank Dr. Don Gibbons for advice on the analysis of the lung tumor pathology. We thank Dr. Richard Behringer for help during the Covid19 pandemic laboratory shutdown. We thank Dr. Awdhesh Kalia, Dr. Nancy A. Jenkins and Dr. Neal G. Copeland for critical comments on the manuscript. We thank members of the Arur Lab for critical discussions during this study. We thank the University of Texas MD Anderson Cancer Center MD Anderson Cancer Center Department of Genetics Microscope core, MD Anderson Cancer Center Advanced Technology Genomic Core (ATGC) (supported by NCI Grant CA016672) and MD Anderson Cancer Center CPRIT Single Cell Genomics Core (supported by grant RP180684). S.A. is the Andrew Sabin Fellow at University of Texas MD Anderson Cancer Center.

## Declaration of interest

S.A. is the inventor of the phosphorylated DICER1 antibody used in this study; a patent application filed by University of Texas systems is pending.

## Lead contact and materials availability

Further information and requests for resources and reagents should be directed to and will be fulfilled by the lead contact, Swathi Arur. This study did not generate new unique reagents.

## Data and code availability

Single-cell RNA-sequencing, ATAC-sequencing and miRNA Nanostring nCounter array data will be deposited at Gene Expression Omnibus (GEO). Any other information and data reported in this paper is available from the lead contact upon request.

## Online Methods

### Nomenclatures used in the study

*KRAS* for human gene; *Kras* for mouse gene; and KRAS for mouse and human protein.

*Dicer1* for mouse gene; and DICER1 for mouse and human protein.

*GKN2* for human gene; *Gkn2* for mouse gene; GKN2 for mouse and human protein.

### Experimental model and subject details

#### Mice (*Mus musculus*) breeding, maintenance, screening, and genotyping

Genotypes of mice used in the study: *Dicer1*^*S2D/+ 12*^, *Kras*^*LA1/+ 8*^ *and Kras*^*LA1/+*^*;Dicer1*^*S2D/+12*^. Mice were maintained in >94% C57BL/J6 background. All mouse studies were conducted in compliance with the institutional animal care and use committee protocol (IACUC). *Dicer1*^*S2D/+*^ animals were crossed to *Kras*^*LA1/+*^ animals to generate a cohort of *Dicer1*^*S2D/+*^, *Kras*^*LA1/+*^ and *Kras*^*LA1/+*^*;Dicer1*^*S2D/+*^ animals. Pups were weaned at 3 weeks of age, ear tagged, and tail snipped for genotyping. Tails were digested in lysis buffer (1M Tris pH 8.0, 5M NaCl, 0.5M EDTA pH 8.0 and 10%SDS) with Proteinase K (20mg/mL) at 55°C overnight. The mice were genotyped each time through tail DNA purification and PCR-based Sanger sequencing. PCR primers used to perform the genotype analysis are presented in the supplementary information. Tumor watch, histopathology and other tumor analyses were performed at indicated timepoints. Moribund animals were euthanized, and their tissues collected for pathology. Animals that were euthanized due to non-tumor related issues (fighting wounds, dermatitis, and others) were not included in the analyses. The number of mice analyzed is stated in each figure. Unless specified, mice of both genders were used. No power analysis was used to determine the sample size.

#### Human lung adenocarcinoma tumor tissue microarrays

Tumor microarrays (TMA) with de-identified patient information were obtained from Discovery Life Sciences (Powell, OH) and TriStar Technology Group, LLC (Washington, DC). 88 lung adenocarcinoma (LUAD) samples from untreated patients were used. No patient or human subjects were recruited for this study. Samples were excluded if they were damaged during sectioning and embedding, core was missing more than 50% of the tissue, key tumor histopathology information was missing or *KRAS* mutation profile was missing. Investigators were blinded for mutation profile and patient clinicopathology information during marker quantification and classification. De-identified patient information used in this paper will be shared by the lead contact upon request.

#### Immunohistochemistry of human tumors

Tumor microarrays (TMA) with 88 lung adenocarcinoma (LUAD) samples from untreated patients were used. Immunohistochemistry (IHC) was performed using a monoclonal anti phospho-DICER1 (1:200, ^13, 14^, and rabbit anti-phospho-ERK (1:100, Thr 202/Tyr 204, Cell Signaling #9101S), diluted in 30% Normal goat serum (NGS) (ThermoFisher, 16210-064). Anti phospho-DICER1 antibody were generated in house and tested for specificity as previously described ^14^. First slides were baked for 1 hour at 60°C, deparaffinized by immersing in Histo-clear solution (National diagnostics, HS-200) for 5 minutes twice, followed by a descending gradient of alcohol washes. Antigen retrieval was performed using 1X Sodium Citrate buffer (Citric acid BDH Cat No. 10081, NaOH BDH Cat No. 30167) by HIER in 2100 Retriever machine (EM Sciences #62700-100). Sections were blocked in 30% Normal goat serum (NGS) for 1 hour at room temperature. After primary antibody incubation, sections were incubated in 1% Hydrogen Peroxide (H_2_O_2_) in methanol for endogenous peroxidase blocking and then in MACH2 Universal polymer-HRP (#M2U522H, BioCare Medical, LLC). Signal was developed with 3,3’-diaminobenzidine (DAB) (Sigmafast™ 3-3’ Diaminobenzidine tablets, D4293-50SET, Sigma). Images were taken using Nikon Eclipse Ti2 equipped with Nikon DS-Ri2 color camera at MD Anderson Cancer Center, Department of Genetics Microscope core. Staining for each antibody was quantified in Fiji following Andy’s Algorithm ^50^ and recorded as a percentage of cells showing positive staining for each protein.

#### Immunofluorescence staining on tissue sections obtained from mice

Tumor tissues and indicated organs analyzed were harvested and fixed in 10% neutral buffered formalin. Right heart ventricle was injected with phosphate buffered saline (PBS, pH 7.4) to perfuse the tissues. Lungs were inflated using formalin through a cannulated trachea. Fixed tissues were paraffin-embedded by the MD Anderson Department of Veterinary Medicine & Surgery Histology Laboratory. Five micrometer sections were used for hematoxylin and eosin (H&E) staining for each of the histological analysis presented. Unstained sections were used for immunofluorescence staining. First, slides were baked for 1 hour at 60°C, deparaffinized by immersing in Histo-clear solution (National diagnostics, HS-200) for 5 minutes twice, followed by a descending gradient of alcohol washes. Antigen retrieved by HIER as described above. Sections were blocked in 30% NGS plus goat anti-mouse IgG (AffiniPure Fab Fragment Goat anti-Mouse IgG, Jackson Immuno Research #115-007-003) for 1 hour at room temperature. The following primary antibodies were diluted in 30% NGS and incubated overnight at 4°C: mouse anti-E-cadherin (1:100, BD #610182), rabbit anti-proSFPC (1:250, Millipore AB3789), guinea pig anti-Vimentin (1:50, Progen, #GP59) and rabbit anti-GKN2 (1:1000, abcam, #ab188866). For GKN2 antibody, Tris-EDTA (10mM Tris base [Sigma, T1503], 1mM EDTA [Sigma, E5134], 0.05% Tween 20 [Fisher BP337), Ph 9.0) was used for antigen retrieving. Samples were then washed in PBS with 1% Tween-20 (PBST). The secondary antibodies were used at 1:500 dilution in PBST: Goat anti-mouse Alexa 488 (Invitrogen, #a11001), Donkey anti-rabbit Alexa 594 (Invitrogen #a21207) and Goat anti-guinea pig Cy5 (Abcam #ab102372). When using anti-IgG Alexa 488 antibody, lung tissue autofluorescence was quenched with Vector TrueView (Vector Laboratory, SP-8400) following manufacturer’s instructions. A 1:1000 dilution of DAPI (2 μg/ml) in 1x PBST was used for nuclear co-staining and slides were mounted in Vectashield (Vector Laboratory, #H-100). Images were acquired using Zeiss Axio Imager.M2 equipped with AxioCam MRm camera at 40x magnification (Zeiss, Thornwood, NY). Images for GKN2 staining were captured on an Evos M7000 (Invitrogen, AMF7000) microscope at 40x magnification.

#### Cell dissociation for single-cell RNA sequencing

Lung tumor cell dissociation was performed following previously published protocol ^51^. Briefly, 39 weeks old mice were anesthetized using Avertin (Sigma, T4802), right heart ventricle was perfused with 1x PBS without calcium and magnesium (ThermoFisher, #100100023). Lungs were removed from the mouse and placed in Liebovitz media (Gibco, #21083-023). Lung tumors were identified and collected under the dissecting microscope. Tumors were minced with forceps and digested in the Liebovitz media with 2 mg/mL collagenase type I (Worthington CLS-1, LS004197), 0.5 mg/mL DNase I (Worthington D LS002007) and 2 mg/mL elastase (Worthington ESL LS002294) for 30 min at 37°C. Digestion was stopped with 20% Fetal bovine serum (FBS, Invitrogen 10082-139). Cells were filtered using a 70 μm cell strainer (Falcon, 352350) on ice and transferred to a 2mL tube. Red blood cells were lysed using Red blood cells lysis buffer (Miltenyl Biotec 130-094-183), and the lysate centrifuged for 1 minute at 5000rpm. Supernatant was removed and cells in the pellet were washed twice with 1mL of ice-cold 1X Leibovitz plus 10% FBS. The cellular suspension was then filtered with a 40 μm cell strainer (Falcon #352340), centrifuged and cells resuspended in ice-cold 1X Leibovitz plus 10% FBS. Cell viability was evaluated using Trypan blue and counted with a hemocytometer. Samples with more than 70% of viable cells were submitted for scRNA-seq at MD Anderson Cancer Center CPRIT Single Cell Genomics Core.

#### Single-cell RNA sequencing and bioinformatic analysis

After cell dissociation and counting, samples were centrifuged and re-suspended in 1X PBS with 0.04% BSA for single cell suspensions and processed for Chromium Single Cells Gene Expression Solution Platform (10x Chromium) with 3’v3 Library following manufacture instructions. Samples were sequence with Illumina NovaSeq6000 S1 with a 2 × 150 paired end configuration and a 400-500 million reads per sample.

FASTQC files were processed using Cellranger v3.1.0 software and mapped to the mouse genome mm10-3.0.0 as reference. Unique molecular identifier (UMI) counts were further analyzed in R v4.1.0 using Seurat v4.0.5 package. Low-quality cells were filtered on the basis of UMI counts (no less than 300), number of transcripts (no more 5000) and mitochondrial transcript fractions (up to 15%). UMI counts were normalized, and scaled using NormalizeData, ScaleData and FindVariableFeatures functions. Principle component analysis was performed, and the number of significant dimensions was estimated using ElbowPlot function. Cell clusters were identified using a shared nearest neighbor (SNN) based algorithm using the FindNeighbors and FindClusters functions. UMAP rendering was performed to visualize clusters. Major cell classes were identified on the basis of enrichment for *Ptprc* (immune cells); *Col1a1, Col3a1*, or *Cdh5* (mesenchymal cells, including endothelial cells); and *Epcam, Cdh1, Sfta2, Lamp3, Rtkn2* (epithelial cells). In addition, automatic annotation with scCATCH v 2.1 package was performed to validate identified cells type. A cell that belong to a major class was subset using the subset function. Each subset was re-clustered and specific cell types were identified on the basis of gene expression using the FindAllMarkers functions. Only genes expressed in at least 25% of the cells per cluster and with a threshold of a log fold change of 0.25 were evaluated. Clusters were visualized using UMAP, as described above. The epithelial cells were identified and annotated based on the gene expression of *Sfta2, Lamp3, Scl34a2, Rtkn2, Pdpn, Hopx, Ager, Foxj1, Scgb1a1* and *Muc5ac*. In addition, heatmaps, volcano plots and feature plots were generated using the DoHeatMap, EnhancedVolcano v1.10.0 and FeaturePlot functions and packages. *Kras*^*LA1/+*^ (K, n=2), *Kras*^*LA1/+*^*;Dicer1*^*S2D/+*^ (KD, n=2) and *Dicer1*^*S2D/+*^ (D, n=1) mice were processed in parallel experimentally and computationally hence they are spatially comparable in UMAP plots.

#### Hairpin Chain Reaction RNA-Fluorescent *in situ* hybridization (FISH)

HCR-RNA-FISH was performed following molecular instruments FFPE tissue sections protocol with minor modifications ^52^. 20 RNA probes were used per gene being analyzed. Slides were baked for 1 hour at 60°C, deparaffinized by immersing in Histo-clear solution (National diagnostics, HS-200) for 5 minutes twice, followed by a descending gradient of alcohol washes. Slides were incubated with 1x Tris-EDTA buffer (pH 9.0) in the 2100 retriever machine for 30 minutes. Then, samples were cooled to 45°C by adding water every 5 minutes. This step was completed in 20 minutes total time. Slides were washed in water, followed by 1x PBS rinse. Proteinase K step was eliminated from the recommended protocol to maintain higher tissue morphology and integrity. The signal was developed by incubating the samples in 200 μL of HCR hybridization buffer inside the humidified chamber at 37°C for 10 minutes, followed by 0.4 pmol of mouse Gkn2-B3 probe and 0.4 pmol of mouse proSfpc-B1 probe in 100uL of HCR hybridization buffer at 37°C overnight. For human tumors, 0.4 pmol of human GKN2-B3 probe and 0.4 pmol of human SFPC-B1 probe was used. Slides were washed in a gradient of HCR probe wash buffer plus 5x SSCT solution (Sigma, S6639). For amplification, slides were incubated with 200 μL of HCR amplification buffer for 30 minutes at room temperature. 100 μL of HCR amplification buffer with 6 pmol of snap-cooled hairpin B3-h1-594, hairpin B3-h2-594, hairpin B1-h1-647, hairpin B1-h2-647 each and DAPI (2 μg/ml) was then added on top of the tissue and incubated overnight in dark humidified chamber at room temperature. Finally, excess hairpins were removed with multiple 5x SSCT washes at room temperature. Slides were mounted with Vectashield (Vector Laboratory, #H-100) and imaged in EvosM700 as described in immunostaining quantification section above.

#### ATAC-sequencing analysis

ATAC-sequencing and bioinformatic analysis was performed by Active motive, Inc. (Carlsbad, CA, USA). A total of 40-50 mg of lung tumors were dissected from euthanized mice, snap frozen in liquid nitrogen and submitted to Active Motif for ATAC-sequencing. At Active motif, samples were processed and put through paired end 42 bp sequencing reads format using Illumina NovaSeq 6000 platform. Reads were mapped to mouse mm10 reference genome using BWA (v0.7.12) algorithm and saved as BAM files. Reads that passed Illumina’s purity filter, aligned with no more than 2 mismatches, and mapped uniquely to the genome were used for further analysis. In addition, duplicate reads (“PCR duplicates”) were removed. For comparative analysis, normalization was achieved by reducing the number of alignments per sample to match the number of alignments in the sample with fewer number of alignments. Reduction was done through random sampling. Genomic regions with a high level of transposition/tagging events were determined using the MACS3 (v3.0.0) peak calling algorithm ^53^, with a cutoff of p-value 1e-7. To identify the density of transposition events along the genome (signal used to generate histograms), the genome was divided into 32 bp bins and the number of fragments in each bin was determined. The histograms showing the density of each signal (peaks) was stored in a bigWig file and visualized using the USCC Genome browser. BigWig files were generated using deeptools (v3.5.1). A FRIP (fraction of reads in peaks) value of 10% or higher was used to define good data quality and false peaks were defined within the ENCODE blacklist ^54^ and removed from analysis. Genomic regions identified by MACS3 in each individual sample were marked by an Interval bar (orange bars below each sample histogram). Finally, because the overlapping Intervals (orange bars) do not have the same length, to be able to compare genomic regions between samples, intervals from all the samples were collected and grouped into “Merged Regions” (green bars on top of histograms). The “Merged Regions” defined the start coordinate of the most upstream region and the end coordinate of the most downstream region. After defining the Intervals and Merged Regions, their peak fragment density, their genomic locations along with their proximities to genes or other genomic features were presented in Excel Spreadsheets (Supplementary information).

### Quantification and statistics analysis

Image quantifications are described in methods for individual experiments. Student t test, chi-square tests and any other statistical analysis of data was done using GraphPad Prism v.9.2.0 software, unless otherwise noted. P values less than 0.05 were considered statistically significant. Scatter or bar plot graphed in GraphPad Prism displayed the SD from the mean value. At least three mice per genotype were used in each experiment, including males and females, unless otherwise noted. Criteria for exclusion are described in experimental model and subject details. The “n” values and statistical test for each experiment are denoted in the corresponding figure panels or graphs.

## Notes

### Competing Interest Statement

The authors have declared no competing interest.

