## Supplementary data for "ERK-dependent DICER1 phosphorylation promotes open chromatin state and lineage plasticity to mediate tumor progression"

### Supplementary methods

#### Quantification of immunofluorescence to assess epithelial-to-mesenchymal transition

Immunofluorescence images for E-cadherin, Vimentin and pro-SFPC stained lungs were captured using Zeiss Axio Imager.M2 with a 40x objective. Prior to image acquisition at 40x, whole lung sections were outlined and mapped with the 10x objective. Tumor regions were identified and marked for further analysis using the Axio Vision Rel 4.8 software. The images of the tumors were acquired upon zooming in on the tissue with the 40x oil objective. At least 5 tumors were analyzed per lung and 5 mice per genotype were used. The total number of cells per optical field was analyzed using FIJI by quantifying the number of DAPI nuclear stained cells, and each cell that expressed the marker being assessed was counted as a positive. Once all the cells in a given optical field were counted, the positive cells were recorded as percentage of total cells per optic field. For GKN2 staining, a similar analysis was carried using the Evos M7000 (Invitrogen, AMF7000), except that the whole lung sections were outlined and mapped with the 2x objective rather than 10x and Evos M7000 software was used to capture the images instead of Axio Imager.

#### miRNA profiling through Nanostring nCounter

For miRNA profiling, 30 to 37 weeks old mice were euthanized following IACUC protocol. Lungs were removed from the mouse, the tumors identified and collected under the dissecting microscope. 50ug of tumors were homogenized in 1 mL of Trizol (Ambion, #15596018). Total RNA was purified following miRNeasy Mini Kit (Qiagen, #217004). 100ng of total RNA was used for Nanostring nCounter miRNA (NS_M_MIR_V1.5) assay, performed by MD Anderson Cancer Center Advanced Technology Genomic Core (ATGC). Results were analyzed in nSolver Analysis Software v4.0. Counts were normalized using internal negative controls, positive controls and the top hundred miRNAs. To calculate ratios and fold change, samples were grouped and partitioned by genotypes. Statistical analysis was performed using two-tailed t-test and Benjamin-Yekutieli for false discovery rate.

#### ChIP-sequencing

ChIP-sequencing and bioinformatic analysis was performed by Active motive, Inc. (Carlsbad, CA, USA). A total of 30 µg of lung tissue from a 20 weeks of age homozygous *Dicer1*$^{S2D/S2D}$ mouse was collected, snap frozen in liquid nitrogen and submitted to Active Motif for Chip-sequencing with anti- DICER1 antibody (Abcam, cat# ab14601, Lot# GR268045-87). Isolated DNA was processed into a standard Illumina Chip-sequencing library and sequenced to generate more than 5 million reads. Reads were aligned to mouse mm10 reference genome using BWA algorithm. Duplicate reads and non-uniquely mapped reads were removed from further analysis. A signal map capturing fragment density along the genome was generated and visualized into Integrated Genome Browser (IGB). Genomic regions with a high signal were determined using the MACS3 (v3.0.0) peak calling algorithm[1], with a cutoff of p-value 1e-7. False peaks were defined using the ENCODE blacklist[2] and removed from analysis.

**Oligonucleotides for PCR and Sanger sequencing genotyping**

| Oligonucleotides |
|---|
| LAI allele genotyping: Lat WT TGCACAGCTTAGTGAGACCC |
| LAI allele genotyping: Lat WT GACTGCTCTCTTTCACCTCC |
| LAI allele genotyping: Lat MUT GGAGCAAAGCTGCTATTGGC |
| Dicer1 S2D genotyping Forward: GGTTTGAGCAGCTTTCGTTAG |
| DICER1 S2D genotyping Reverse: CCCTGTCACTGAAACATGAAC |
| KRas G12D sequencing Forward: GCCTGCTGAAAATGACTGAGTATAA |
| KRas G12D sequencing Reverse: AGGGTCATACTCATCCACAAAGT |
| DcrS1712D_Blast_F: GCGCAGATTGTTACCAGCG |
| DcrS1712D_Blast_R: CAACTATCCCTCTGGACAGCC |
| DcrS1836D_Blast_F: TCTGGCAGGTGTACTATCCGA |
| DcrS1836D_Blast_R: AACGTGGACGCTGAGAGGAT |

### Supplemental figure titles and legends

**Supplementary Table 1.** Patient clinicopathology features between *KRAS* and non-*KRAS* mutated LUADs.

| **Patient characteristics** | ***KRAS* (n= 38)** | | | **Non-*KRAS* (n= 50)** | | |
|---|---|---|---|---|---|---|
| | **Median** | **Min** | **Max** | **Median** | **Min** | **Max** |
| **Age** | 71 | 41 | 85 | 73 | 42 | 87 |
| | **N (%)** | | | **N (%)** | | |
| **Sex** | | | | | | |
| **Female** | 21 (55%) | | | 25 (50%) | | |
| **Male** | 16 (42%) | | | 24 (48%) | | |
| **Undocumented** | 1 (3%) | | | 1 (2%) | | |
| **Stage** | | | | | | |
| **I** | 15 (39%) | | | 29 (58%) | | |

| | | |
|---|---|---|
| **II** | 9 (23%) | 14 (28%) |
| **III** | 5 (13%) | 5 (10%) |
| **IV** | 0 (0%) | 0 (0%) |
| **Sx** | 9 (23%) | 2 (4%) |
| **Tumor** | | |
| **T1** | 15 (39%) | 11 (22%) |
| **T2** | 15 (39%) | 25 (50%) |
| **T3** | 5 (13%) | 11 (22%) |
| **T4** | 0 (0%) | 1 (2%) |
| **Tx** | 3 (8%) | 2 (4%) |
| **Node** | | |
| **N0** | 20 (53%) | 40 (80%) |
| **N1** | 9 (23%) | 8 (16%) |
| **N2** | 4 (11%) | 1 (2%) |
| **N3** | 0 (0%) | 0 (0%) |
| **Nx** | 5 (13%) | 1 (2%) |

Note: Tx, Sx or Nx, show samples were patient information was not evaluated or undocumented at time of sample collection.

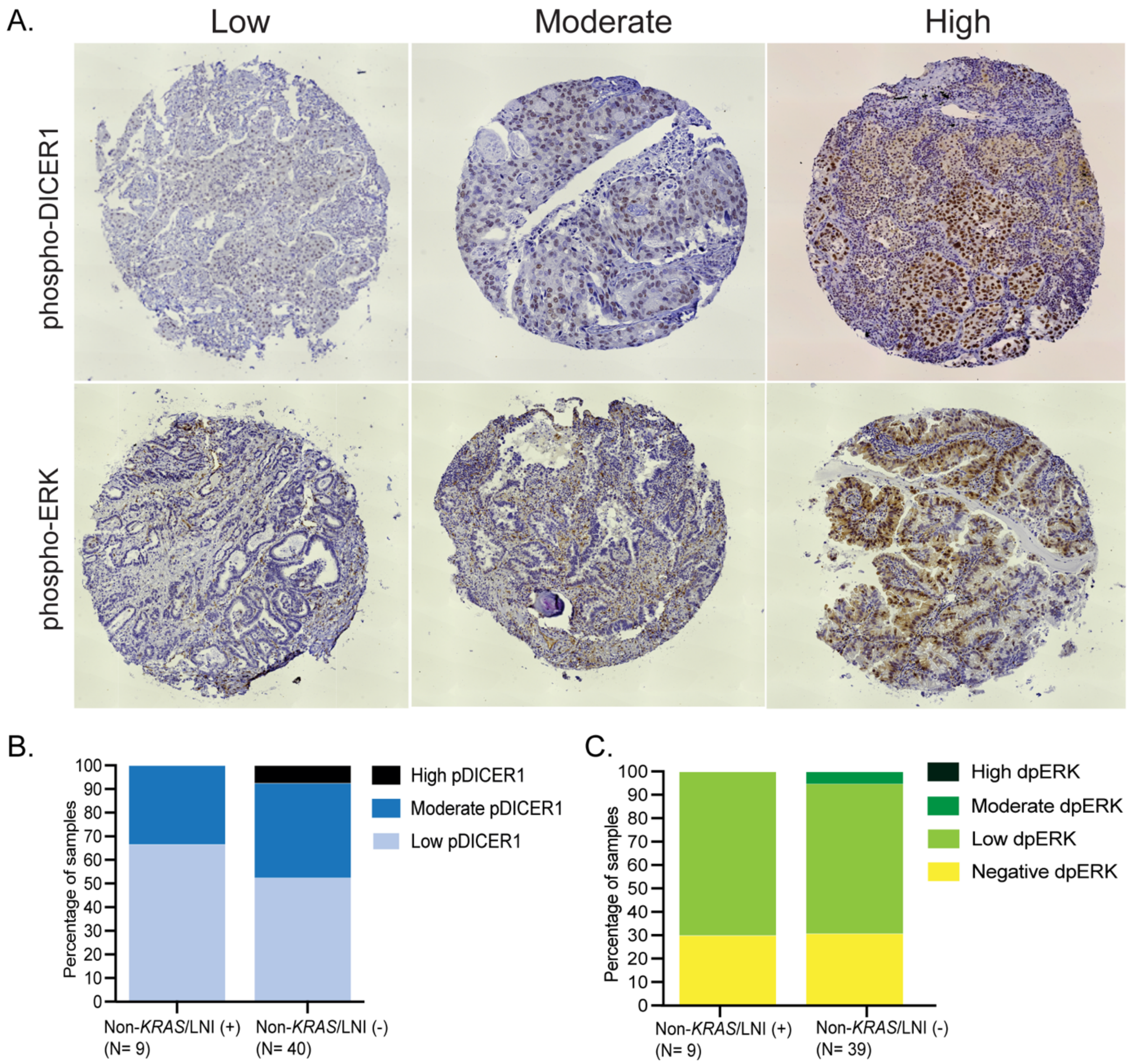


**Supplementary Figure 1. Phosphorylated DICER1 expression in human LUAD tumors**

A. Representative images of human LUAD tumors stained with anti phospho-DICER1 and anti phospho-ERK antibodies. Samples are plotted by percentage of cells positive for phospho-DICER1 in brown (top panel), or phospho-ERK (bottom panel). Negative (<10% positive cells), Low, (≥10% <30% positive cells), moderate (≥30% < 70% of positive cells), high (≥70% of positive cells). 40x.
B. Human LUAD tumors with no mutations in *KRAS* are plotted based on presence (LNI (+)) or absence (LNI (-)) of lymph node invasion and phospho-DICER1 positive signal. Chi-square, p-value= 0.2058.
C. Human LUAD tumors with no mutations in *KRAS* are plotted based on presence (LNI (+)) or absence (LNI (-)) of lymph node invasion and phospho-ERK positive signal. Chi-square, p-value= 0.1048.

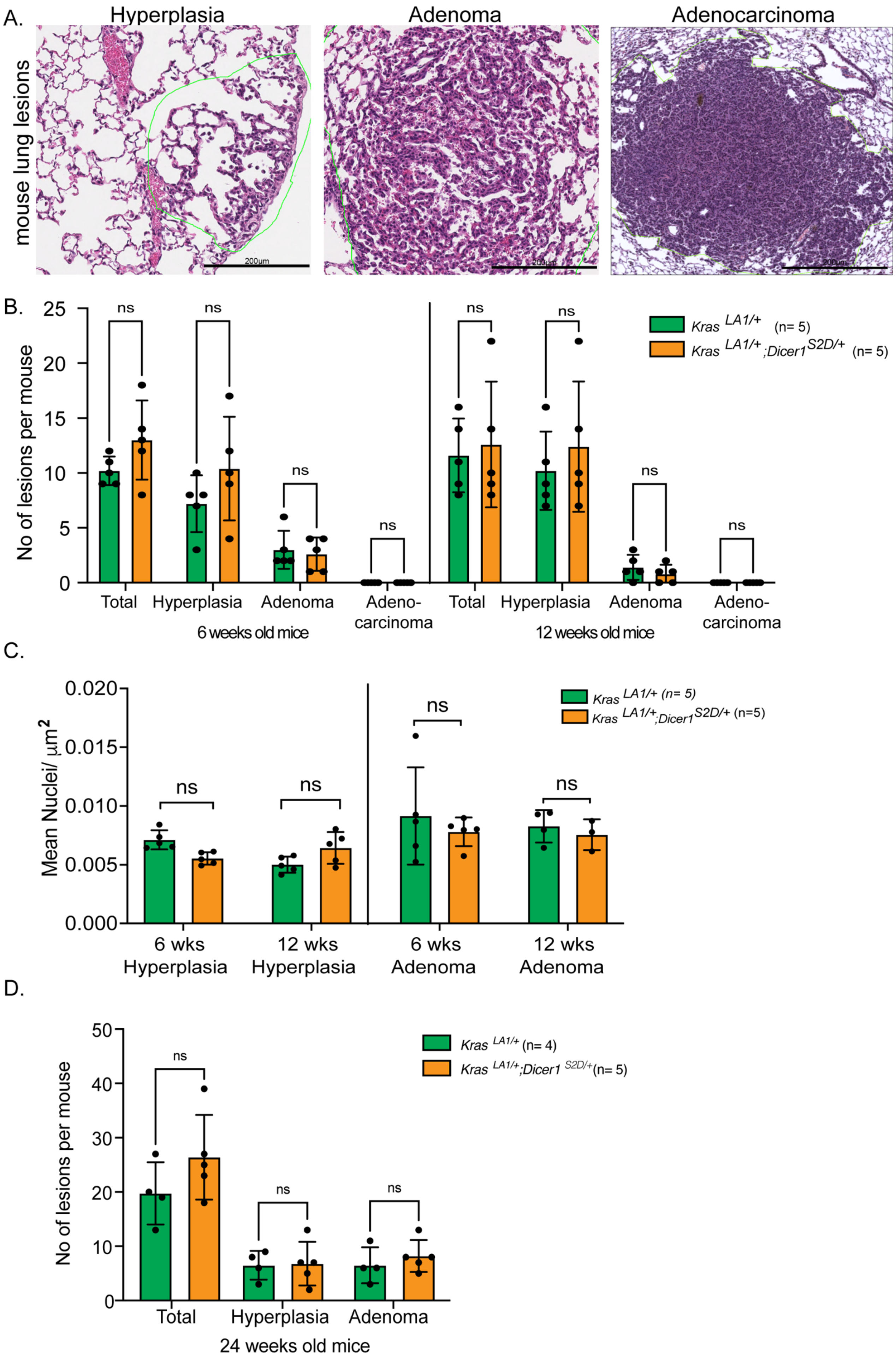
A.
Hyperplasia
Adenoma
Adenocarcinoma
mouse lung lesions
200µm
B.
No of lesions per mouse
25
20
15
10
5
0
ns
Total
Hyperplasia
Adenoma
Adeno-carcinoma
6 weeks old mice
12 weeks old mice
Kras LA1/+ (n= 5)
Kras LA1/+;Dicer1 S2D/+ (n= 5)
C.
Mean Nuclei/ µm2
0.020
0.015
0.010
0.005
0.000
ns
6 wks Hyperplasia
12 wks Hyperplasia
6 wks Adenoma
12 wks Adenoma
Kras LA1/+ (n= 5)
Kras LA1/+;Dicer1 S2D/+ (n=5)
D.
No of lesions per mouse
50
40
30
20
10
0
ns
Total
Hyperplasia
Adenoma
24 weeks old mice
Kras LA1/+ (n= 4)
Kras LA1/+;Dicer1 S2D/+ (n= 5)

**Supplementary Figure 2. Phosphomimetic *Dicer1* leads to late-stage of tumor progression**

A. Representative images of lung lesions and their classification by nomenclature. Scale bar = 200μm
B. Number of lung lesions in mice from *Kras*$^{LA1/+}$ (n= 5) in green and *Kras*$^{LA1/+}$;*Dicer1*$^{S2D/+}$ (n= 5) in orange at 6 and 12 weeks of age. p-value > 0.05.
C. Tumor size quantified as total number of cells divided by the tumor area for *Kras*$^{LA1/+}$ (n= 5) in green and *Kras*$^{LA1/+}$;*Dicer1*$^{S2D/+}$ mice (n= 5) in orange at 6 and 12 weeks of age. p-value > 0.05.
D. Number of lung lesions in mice from *Kras*$^{LA1/+}$ (n= 4) in green and *Kras*$^{LA1/+}$;*Dicer1*$^{S2D/+}$ (n= 5) in orange at 24 weeks of age. Number of adenocarcinomas in main figure 2. p-value > 0.05.

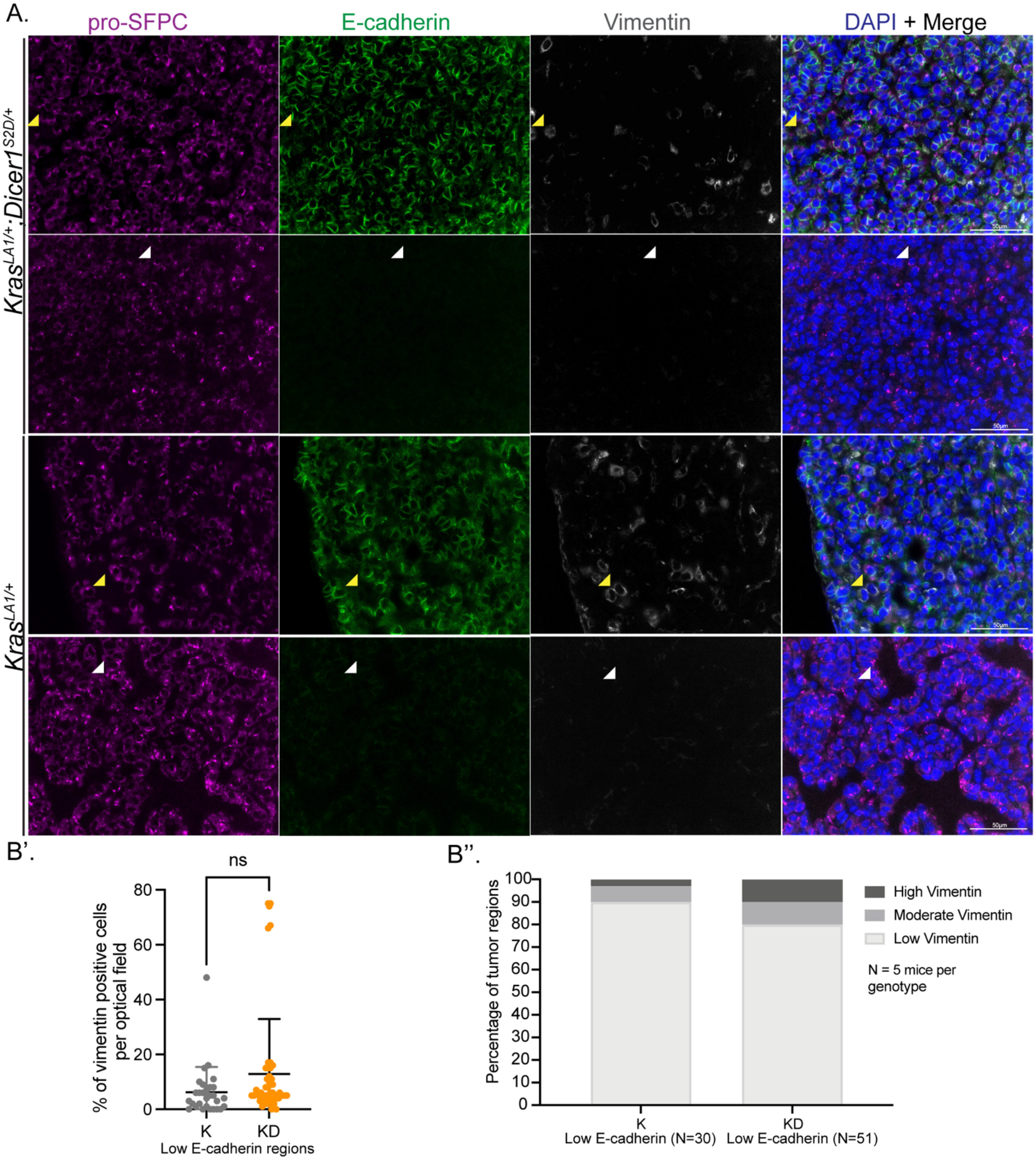


**Supplementary Figure 3. Phosphomimetic *Dicer1* does not affect EMT in lung tumors**

A. Representative images of lung tumors from *Kras*$^{LA1/+}$ (n=5) and *Kras*$^{LA1/+}$;*Dicer1*$^{S2D/+}$ (n=5) at ~52 weeks of age stained with pro-surfactant C (pro-SFPC), E-cadherin and Vimentin antibodies. Scale bar = 50μm. Yellow arrowhead points to Vimentin positive cells. White arrowhead points to AT2 cells that have low E-cadherin and no Vimentin signal.

B. B' Scatter plot showing the percentage of vimentin positive cells in areas of the tumors where AT2 cells showed low E-cadherin staining in *Kras*$^{LA1/+}$ (K, n=5) and *Kras*$^{LA1/+}$*;Dicer1*$^{S2D/+}$ (KD, n=5) animals at 52 weeks of age. Quantification was done per optical field in a 40X magnification (Methods).

B'' Bar graphs showing the distribution of tumor regions that showed Vimentin positive cells in areas where AT2 cells showed low E-cadherin staining. Optical fields were classified as Low, (<30% of Vimentin positive cells), moderate (≥30% < 70% of Vimentin positive cells), high (≥70% of Vmentin positive cells).

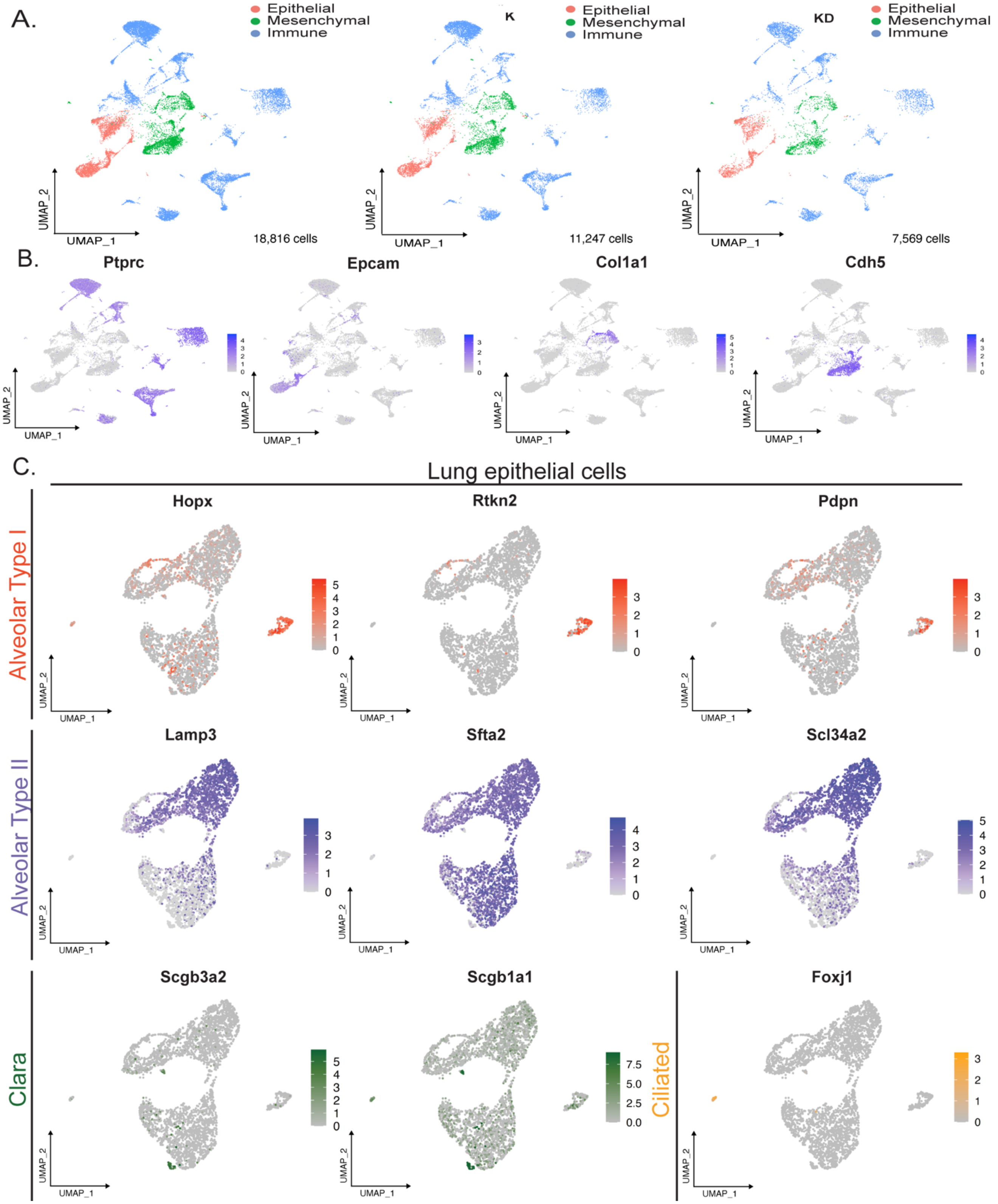

**Supplementary Figure 4. scRNA-Seq analysis of *Kras*$^{LA1/+}$ and *Kras*$^{LA1/+;}$*Dicer1*$^{S2D/+}$ lung tumors**

A. UMAP graphs of the distinct cell types identified from single-cell RNA sequencing analysis of lung tumors of *Kras*$^{LA1/+}$ (K, n=2) and *Kras*$^{LA1/+}$*;Dicer1*$^{S2D/+}$ (KD, n=2) mice. The left panel shows a merge of data from the two genotypes of K and KD.
B. Feature plot showing markers (in purple) used to identify immune cells (*Ptprc*, CD45), mesenchymal (*Col1a1* and *Cdh5*) and epithelial cells (*Epcam*).
C. Feature plots for the gene expression profiles that mark the distinct lung epithelial cell types. Alveolar type I (AT1) in red, Alveolar type II (AT2) in purple, Clara in green and Ciliated cells in yellow.

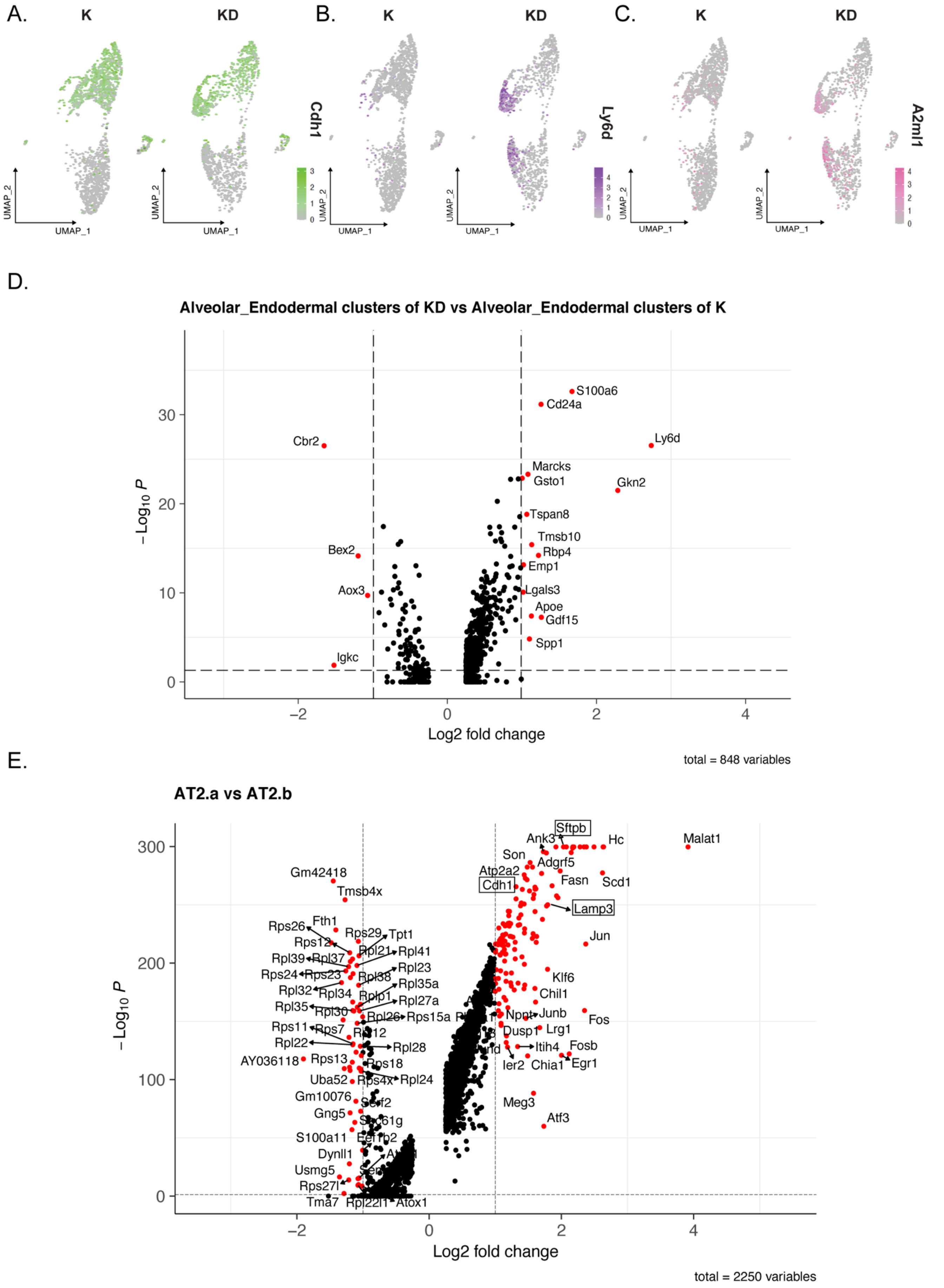


**Supplementary Figure 5. Phosphomimetic *Dicer1* reprograms a subset of alveolar type II tumor cells**

A. UMAP graph showing E-cadherin (*Cdh1*) gene expression in green in epithelial tumor cell clusters from *Kras*$^{LA1/+}$ (K, n=2) and *Kras*$^{LA1/+}$*;Dicer1*$^{S2D/+}$ (KD, n=2) animals.
B. UMAP graph showing *Ly6d* gene expression, an esophageal gene, in purple in the epithelial tumor cell clusters from *Kras*$^{LA1/+}$ (K, n=2) and *Kras*$^{LA1/+}$*;Dicer1*$^{S2D/+}$ (KD, n=2) mice.
C. UMAP graph showing *A2ml1* gene expression in pink in tumor epithelial clusters of *Kras*$^{LA1/+}$ (K, n=2) and *Kras*$^{LA1/+}$*;Dicer1*$^{S2D/+}$ (KD, n=2) mice.
D. Volcano plot showing significantly upregulated and downregulated genes (red dots) in the alveolar_endodermal cluster of *Kras*$^{LA1/+}$*;Dicer1*$^{S2D/+}$ (KD, n=2) when compared to alveolar_endodermal cluster of *Kras*$^{LA1/+}$ mice.
E. Volcano plot showing significantly upregulated and downregulated genes (red dots) in the AT2.a cluster when compared to AT2.b cluster. Highlighted with squares are *Cdh1*, *Lamp3* and *Sftpb*.

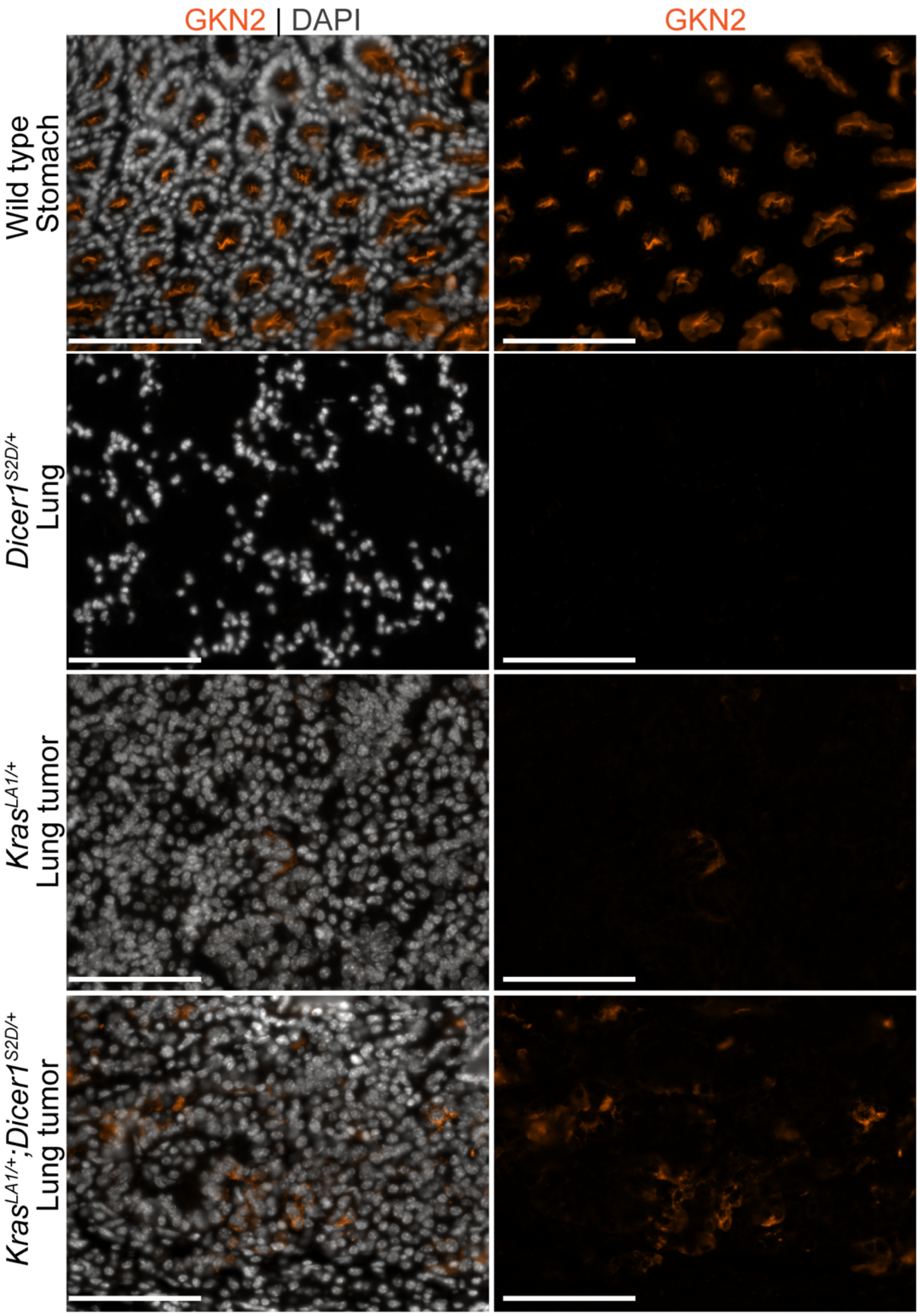


**Supplementary Figure 6. Phosphomimetic *Dicer1* alters the identity of AT2 tumor cells.**

Representative images of *Gkn2* protein expression (in orange) and nuclei (DAPI in grey) in wild type stomach (positive control for *Gkn2*), *Dicer1*$^{S2D/+}$ lung and lung tumors from *Kras*$^{LA1/+}$ and *Kras*$^{LA1/+}$;*Dicer1*$^{S2D/+}$ mice. Scale bar = 50μm

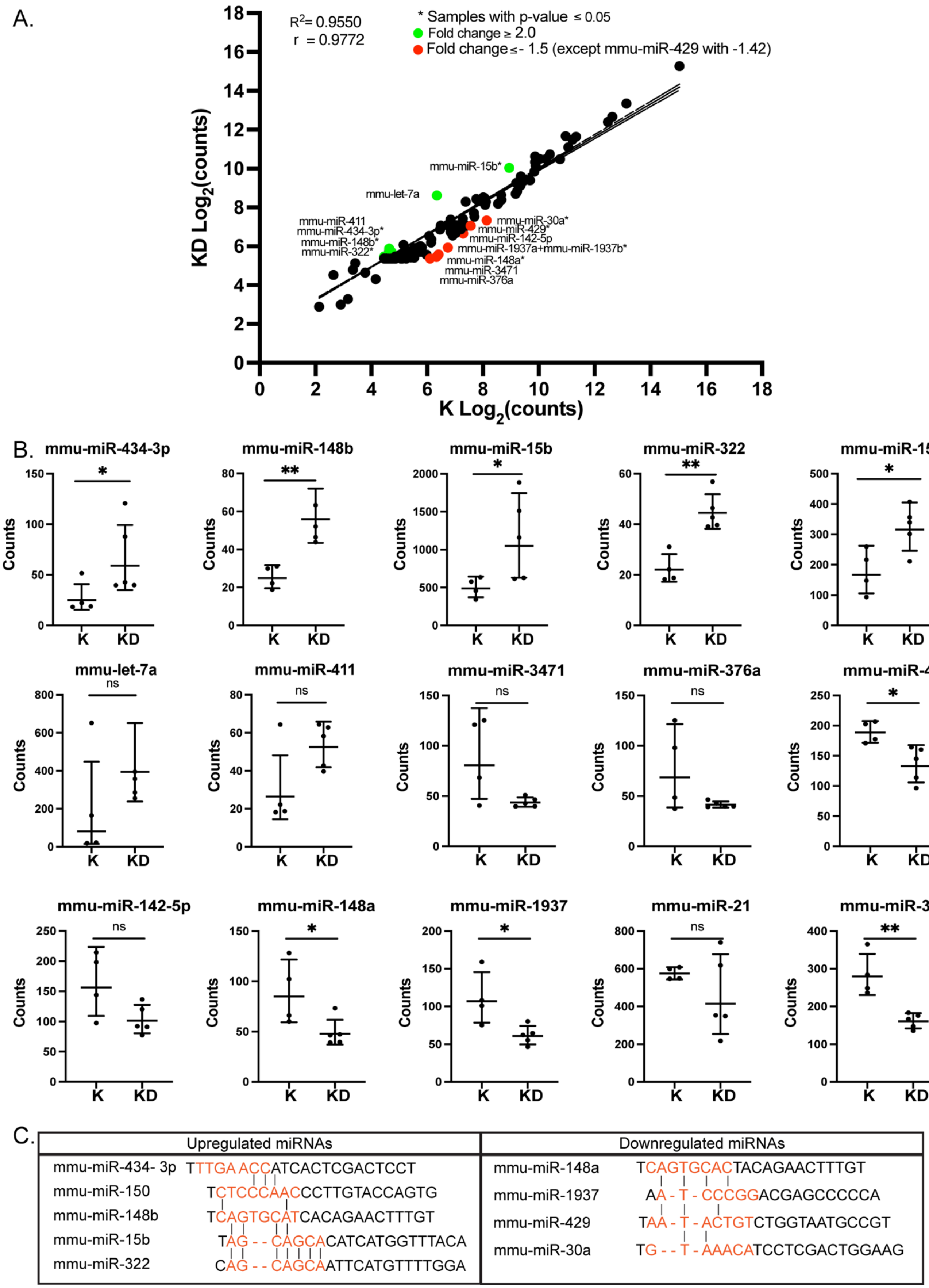


**Supplementary Figure 7. Phosphomimetic *Dicer1* does not significantly affect microRNA production**

A. Graph showing simple linear regression of miRNAs expression as assessed by Nanostring nCounter analysis on $Kras^{LA1/+}$ (K, n=6) lung tumors plotted on the x-axis and $Kras^{LA1/+}$;$Dicer1^{S2D/+}$ (KD, n=6) lung tumors plotted on the y-axis. Each dot represents a mature miRNA. In green are miRNAs with at least a 2.0-fold increase and in red are miRNAs with at least a 1.5-fold decrease.

B. The graphs show individual scatter plots of miRNAs from Panel A that were either upregulated (green) and or downregulated (red).

C. Alignment of mature miRNAs that were significantly upregulated and downregulated in the $Kras^{LA1/+}$;$Dicer1^{S2D/+}$ lung tumors relative to $Kras^{LA1/+}$ lung tumors. Seed sequences are shown in orange.

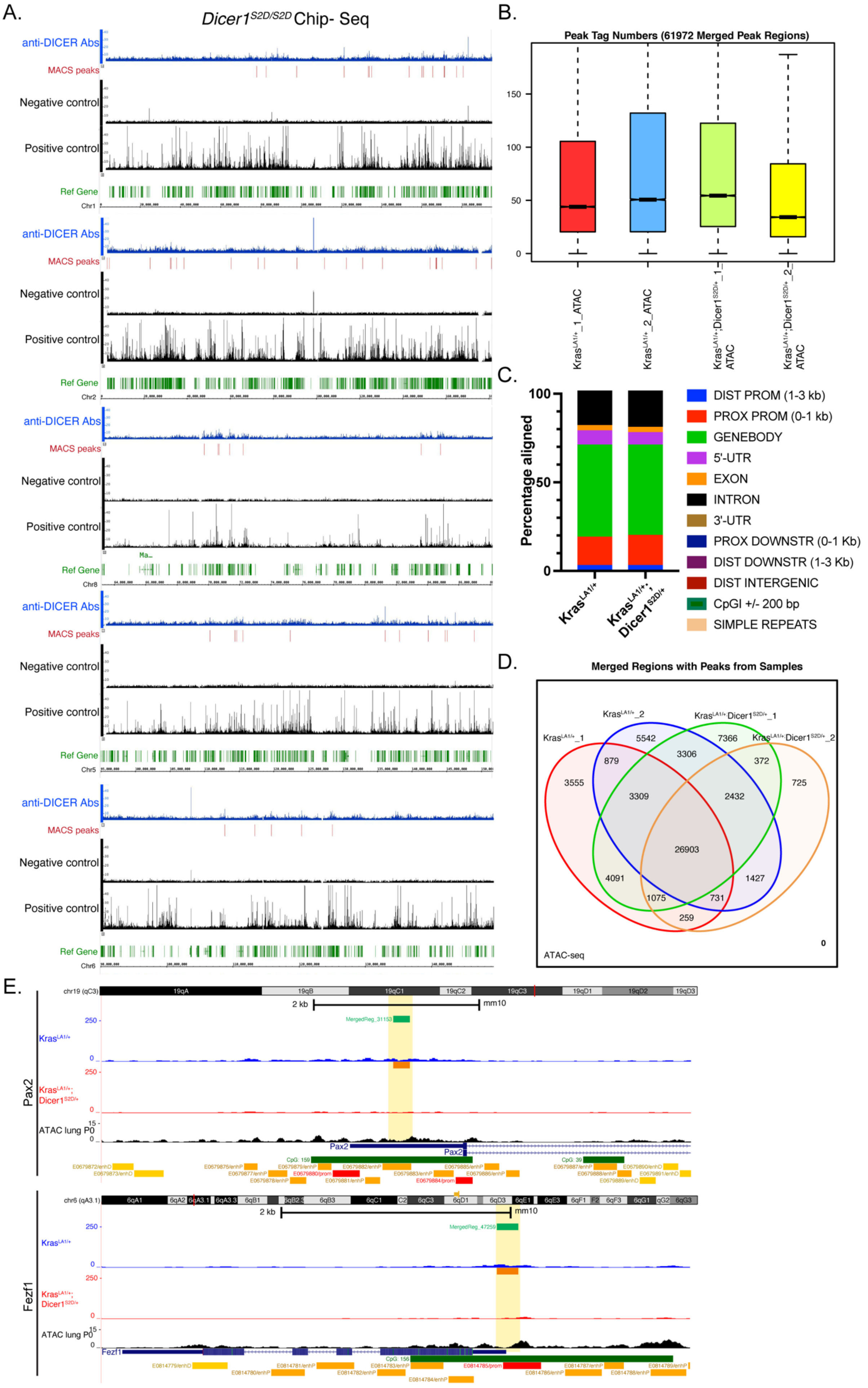
A.
Dicer1S2D/S2D Chip- Seq
anti-DICER Abs
MACS peaks
Negative control
Positive control
Ref Gene
Chr1
Chr2
Chr8
Chr5
Chr6
B.
Peak Tag Numbers (61972 Merged Peak Regions)
0
50
100
150
KrasLA1/+_1_ATAC
KrasLA1/+_2_ATAC
KrasLA1/+;Dicer1S2D/+_1_ATAC
KrasLA1/+;Dicer1S2D/+_2_ATAC
C.
Percentage aligned
100
50
0
KrasLA1/+
KrasLA1/+; Dicer1S2D/+
DIST PROM (1-3 kb)
PROX PROM (0-1 kb)
GENEBODY
5'-UTR
EXON
INTRON
3'-UTR
PROX DOWNSTR (0-1 Kb)
DIST DOWNSTR (1-3 Kb)
DIST INTERGENIC
CpGI +/- 200 bp
SIMPLE REPEATS
D.
Merged Regions with Peaks from Samples
KrasLA1/+_1
KrasLA1/+_2
KrasLA1/+;Dicer1S2D/+_1
KrasLA1/+;Dicer1S2D/+_2
3555
5542
7366
725
879
3306
372
3309
2432
26903
4091
1427
1075
731
259
0
ATAC-seq
E.
Pax2
chr19 (qC3)
2 kb
mm10
MergedReg_31153
KrasLA1/+
KrasLA1/+; Dicer1S2D/+
ATAC lung P0
250
0
15
Fezf1
chr6 (qA3.1)
MergedReg_47259

**Supplementary Figure 8. Phosphomimetic nuclear DICER1 does not directly bind to chromatin to alter its compaction state**

A. Histograms showing low peak signal across multiple chromosomes of *Dicer1*$^{S2D/S2D}$ lung tissues chip-sequencing analysis with anti-total DICER1 antibody (in blue). Black tracks show positive and negative controls retrieved from the database for comparison. Chromosome coordinates and reference genes (in green) are displayed below each histogram. MACS peaks are displayed in red bars.

B. Box plot showing the peak distribution for each sample analyzed in ATAC-sequencing. *Kras*$^{LA1/+}$_1 and *Kras*$^{LA1/+}$_2 represented biological replicates. *Kras*$^{LA1/+}$;*Dicer1*$^{S2D/+}$_1 and *Kras*$^{LA1/+}$;*Dicer1*$^{S2D/+}$_2 represent biological replicates. The boxed area represents the center two quartiles, notched line represents the mean and the whiskers show the top and bottom quartiles without outliers.

C. Distribution of identified peaks based on their genomic alignment in *Kras*$^{LA1/+}$ (n=2) and *Kras*$^{LA1/+}$;*Dicer1*$^{S2D/+}$ mice lung tumors. Each genomic region is color coded as described in the legend.

D. Ven diagram showing the number of Merged Regions that overlap between samples. *Kras*$^{LA1/+}$_1 and *Kras*$^{LA1/+}$_2 represented biological replicates. *Kras*$^{LA1/+}$;*Dicer1*$^{S2D/+}$_1 and *Kras*$^{LA1/+}$;*Dicer1*$^{S2D/+}$_2 represent biological replicates.

E. Representative histogram showing ATAC-sequencing results for *Kras*$^{LA1/+}$ (in blue), *Kras*$^{LA1/+}$; *Dicer1*$^{S2D/+}$ (in red) lung tumors, and wild type mouse lung (in black) at *Pax2* and *Fezf1* locus. ATAC-sequencing from wild type mouse lung was retrieved from ENCODE Consortium 3 database. Green bars show the "Merged Regions" as described in Methods. Orange bars shows individual Intervals for each sample as described in Methods. For each gene, exons are represented as rectangular bars and introns as lines with arrows (pointing the direction of transcription). Regulatory elements in the mouse genome as identified by the ENCODE Registry of candidate cis-Regulatory Elements (cCREs) database are shown in the bottom of each panel. Highlighted in light yellow are regions absent in the *Kras*$^{LA1/+}$;*Dicer1*$^{S2D/+}$ lung tumors and present in the *Kras*$^{LA1/+}$ lung tumors.
